# *Salmonella* effector kinase SteC is activated by host-mediated phosphorylation

**DOI:** 10.1101/2025.03.01.640964

**Authors:** Timesh D. Pillay, Briac Lemetais, Jessica Huber, Ines Diaz del Olmo, Yan Li, Laura Masino, Sarah Maslen, Mei Liu, Diego Esposito, Jay C.D. Hinton, Xiujun Yu, Teresa L.M. Thurston, Katrin Rittinger

**Author notes:** Correspondence (T.L.M.T.), (K.R.).

## Abstract

The pathogen *Salmonella*, which causes significant human morbidity and mortality, encodes an effector kinase, SteC, which mediates actin polymerisation and cell migration. Given the minimal nature of its kinase domain, it remains unclear how SteC is catalytically active and how this activity is regulated. Here, we show that SteC is activated following the phosphorylation of the highly conserved S379 residue by a host kinase. Phosphorylation of S379 dramatically increases nucleotide binding affinity of SteC, enabling substrate phosphorylation and promoting actin polymerisation. Further mutational analysis identified the functional role of HD and DGD motifs that likely mimic the HxD and DFG motifs of eukaryotic kinases. Meanwhile, the C-tail of SteC, encompassing amino acids 429-457, is essential for function following translocation from *Salmonella*, but dispensable for catalysis *in vitro*. Overall, our findings uncover two previously unappreciated mechanisms that mediate the activity of the only *Salmonella* effector kinase within the host.

**HIGHLIGHTS:** SteC is phosphorylated at S379 in mammalian cells

SteC phosphorylation is critical for kinase activity and actin polymerisation

Phosphorylation at S379 promotes nucleotide binding

The C-tail of SteC is not required for catalysis but is essential for activity in infected cells

## INTRODUCTION

The Gram-negative facultative intracellular pathogen, *Salmonella enterica*, causes >100 million human infections globally each year. Disease manifestations include gastroenteritis, invasive non-typhoidal *Salmonella* disease and typhoid fever, that together cause an estimated 242,500 deaths worldwide annually^1–3^. To facilitate their intracellular lifestyle, serovars of *Salmonella enterica* utilise two type 3 secretion systems (T3SSs) to translocate effector proteins from the bacterium into the host cell cytoplasm. Together, these effectors trigger a range of phenotypes that include invasion of non-phagocytic cells, generation and maintenance of the *Salmonella*-containing vacuole (SCV) and inhibition of host cell-intrinsic immunity^4^. The translocated effectors demonstrate a diverse array of biochemical activities, many of which are eukaryote-like^4^. One such activity is the addition of phosphate to target substrate proteins, which represents one of the most abundant post translational modifications found in eukaryotes. Phosphorylation provides precise, reversible, regulation of diverse, core cellular processes including metabolism, growth and cell cycle progression as well as immune signalling and apoptosis. *Salmonella* expresses a single eukaryotic-like serine-threonine (S/T) kinase effector called SteC, which has been reported to phosphorylate mitogen-activated protein kinase, MEK1^5^; heat shock protein, HSP27^6^; the formin-like proteins, FMNL1/2/3^7^; and myosin light chain protein MYL12A^8^. The kinase activity of SteC has been reported to support the generation of a dense meshwork of F-actin surrounding the micro-colony^9^, actin rearrangement and macrophage migration^8^. Despite this, little is known regarding how SteC functions enzymatically and how its activity is regulated.

Eukaryotic kinases have a conserved structural core consisting of an N lobe that binds ATP and a C lobe that binds and orients the substrate towards the catalytic cleft between the two lobes. Several amino acid motifs and residues that are essential for catalysis have been characterised, with canonical kinases consisting of 12 conserved subdomains^10–14^. Based on homology to eukaryotic S/T kinases, SteC retains N lobe features of the glycine-rich loop in subdomain I, which mediates ATP binding and the catalytic invariant lysine (K256) in subdomain II that anchors ATP and catalyses phosphoryl transfer^9^. A glutamic acid in alpha helix (αC) of subdomain III normally stabilises interaction between the invariant lysine in subdomain II and ATP. In SteC, this is proposed to be E272^9^, but this has not been experimentally validated. The structure of the N lobe of SteC has recently been reported (PDB: 8JBI)^8^. The N-lobe crystallises as a dimer leading to a proposed model of activation whereby ATP is stabilised at the dimeric interface of two SteC molecules. However, this portion of the kinase is catalytically inactive^8^, raising questions concerning the biological relevance of the model. Furthermore, the C lobe of SteC is significantly smaller than others found in many eukaryotic kinases. Its minimal size mirrors the *Shigella* effector, OspG, which requires binding to an E2-ubiquitin conjugate to become active^15^. Together, this raises the question as to how SteC kinase activity is controlled after its delivery into host cells.

Here we report that at the mechanistic level, SteC is activated upon phosphorylation of S379 by a host kinase. S379 of SteC resides in a putative activation segment and is required for 1) enhanced nucleotide binding 2) to enable phosphorylation of known substrates, FMNL1 and MYL12A, and 3) to mediate SteC-induced actin polymerisation. Our findings therefore reveal how phosphorylation controls the activation of this bacterial virulence factor.

## RESULTS and DISCUSSION

### SteC represents a minimal kinase with a depleted C lobe

Despite SteC having proposed kinase activity towards multiple substrates, a detailed understanding of its catalytic mechanism remains to be determined. The AlphaFold2 predicted structure of SteC shows an N-terminal regulatory domain and C-terminal kinase domain, connected by a linker (**Figure 1a** and **S1a**). Within the kinase domain, the N-terminal lobe (subdomains I-IV) mediates anchoring and orienting of the ATP molecule and comparison with the canonical eukaryotic kinase PKA (PDB: 1ATP^16^) demonstrates that this is largely conserved in SteC (**Figure 1a** and **S1b**). However, the C lobe, which normally consists of subdomains VIa-XI and mediates substrate binding and initiates phosphate transfer, is notably diminished in SteC. Instead, in SteC, this region spanning amino acids 320-457, is predicted to consist of αE, β6-9 and a further two α-helices (αF and αG) followed by a largely unstructured C-terminal tail (“C-tail”) (amino acids 429-457) (**Figure 1a and S1b**). It appears that H343 and D344 may take on the role of the H*D motif in canonical kinases (**Figure S1b**). It is however unclear whether SteC contains a DFG motif (normally residing between β8 and β9), or an activation segment, which would canonically start at the DFG motif and extend to the APE motif within αEF. The aspartate of the DFG motif mediates interaction with the Mg^2+^ ion to aid positioning of the gamma-phosphate. Whether this function is mediated instead by a DGD motif is unclear (**Figure 1a and S1b**). No APE-like motif, found at the C-terminal end of the activation segment is evident in SteC (**Figure S1b**). Furthermore, the only crystalised fragment of SteC, covering amino acids 202-375, terminates at the end of αEF, leaving the role of the two remaining alpha helices (αF and αG) and unstructured C-tail unknown.

**Figure 1:**
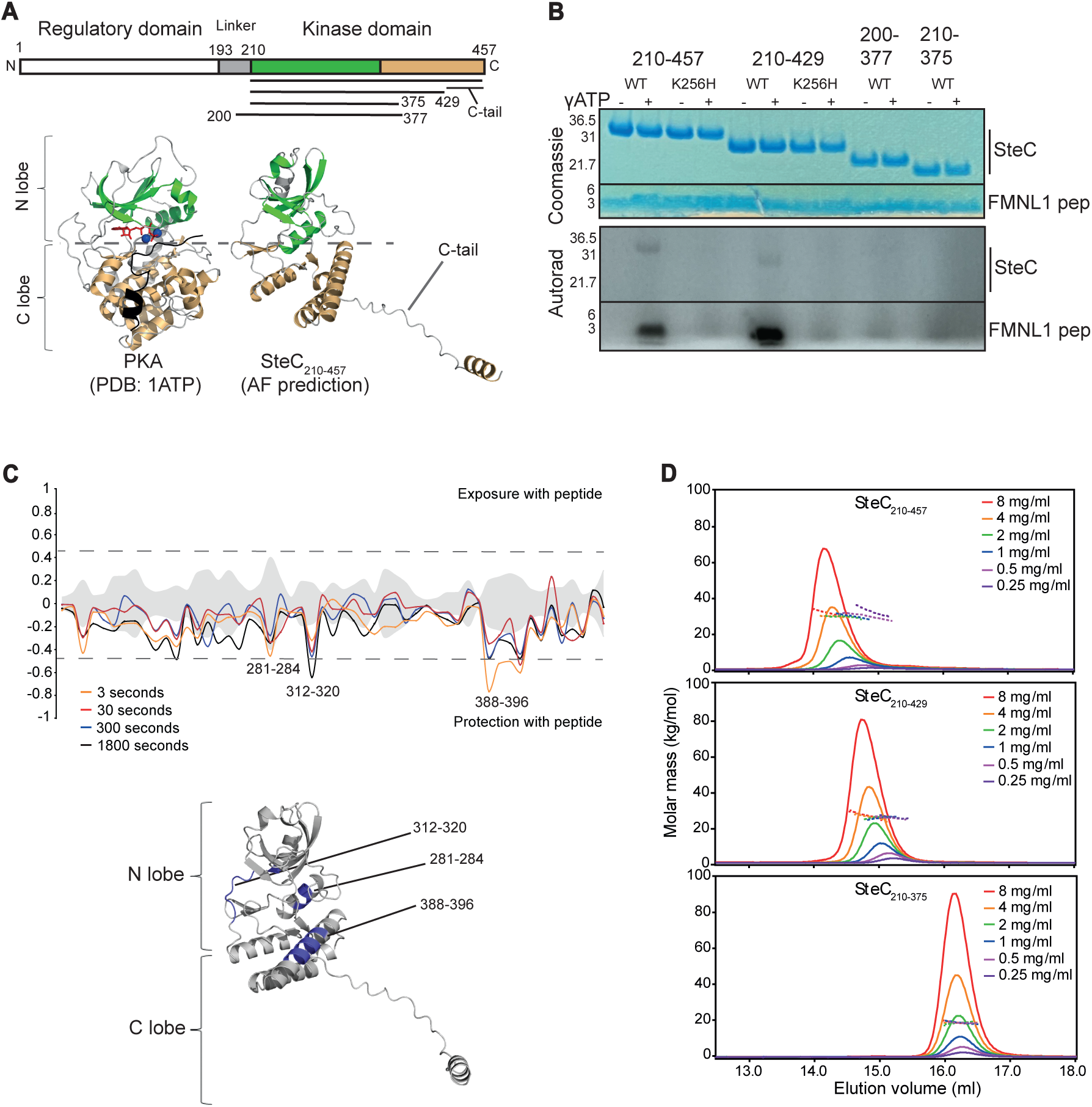
SteC is monomeric in solution. a) Top: Schematic of SteC, showing two domains connected by a linker. The kinase domain contains N lobe (green) and C lobe (light orange). The C-tail (amino acids 430-457) and the three kinase domain constructs used in this study are indicated. Bottom: cartoon representation of PKA (left) in complex with peptide substrate (black), ATP (red) and magnesium (blue) (PDB: 1ATP, left) and SteC kinase domain AF2 prediction (right) in the same orientation. The N lobes (green), C lobes (light orange) and loops (grey) are indicated, as are the catalytic clefts by a dotted line. b) Radioactive kinase assays of 5 μM SteC_210-457_ WT or K256H, SteC_210-429_ WT or K256H, SteC_200-377_ and SteC_210-375_ expressed in *E. coli* with 100 μM FMNL1 peptide. c) HDX analysis of SteC_210-457;K256H_ in complex with FMNL1 peptide. Exchange is plotted as a relative difference in deuterium uptake of SteC_210-457;K256H_ alone or in the presence of the FMNL1 K190 peptide across the amino acid range. See **Figure S1e** for peptide coverage. d) SEC-MALLS analysis of the three SteC kinase domain fragments SteC_210-457_, SteC_210-429_, and SteC_210-375_, expressed with K256H mutation. The dotted lines represent the corresponding estimated molar mass of the eluting species.

### The depleted C lobe of SteC mediates substrate binding and phosphorylation

To investigate the role of the C lobe of SteC, and test whether catalytic residues reside within this region, we expressed and purified three truncated forms of SteC: the entire predicted kinase domain (210-457), the kinase domain lacking the C-tail (210-429) and constructs 200-377 and 210-375 which lack αF and αG as well as the C-tail (**Figure 1a and S1c**). 1D ^1^H-NMR analysis revealed that the smallest construct, SteC_200-377_, produced a spectrum with the sharpest peaks covering a wider range of chemical shifts (**Figure S1d**) indicative of a well folded construct. Although a subset of peaks in the spectra of the longer SteC constructs, SteC_210-457_ and SteC_210-429_, have the same chemical shifts as in the SteC_200-377_ spectrum, they exhibited significant peak broadening. This effect could be attributed to the constructs’ higher molecular weights and the presence of chemical exchange from either inter- or intra-molecular interactions. Interestingly, given the similarity of SteC_210-457_ and SteC_210-429_ spectra, there is little evidence that the additional C-terminal residues are completely flexible. This suggests that contrary to the AlphaFold prediction, the C-tail of SteC might exist in equilibrium between free and bound states, where it interacts intramolecularly with the SteC core (**Figure S1d**).

Next, the ability of SteC_210-375_, SteC_210-429_ and SteC_210-457_ to mediate substrate phosphorylation was analysed by radioactive kinase assays, using the reported substrate FMNL1^7^. Incubation of FMNL1_1-458_ with SteC and ATP resulted in the identification of three phosphorylation sites (S199, S203 and S207) (**Table S1**). As this protein is prone to aggregation, an FMNL1 peptide, spanning amino acids K190-K213 was generated. Whereas SteC_200-377_ and SteC_210-375_ were inactive, SteC_210-457_ and SteC_210-429_ phosphorylated the FMNL1 peptide, and this was dependent on the invariant lysine, K256 (**Figure 1b**). Absence of substrate phosphorylation by SteC_210-375_ and SteC_200-377_ corroborates previous findings^8^ and is unsurprising as this truncated form of SteC lacks a significant portion of the C-lobe which forms part of the substrate binding module. To identify the regions of SteC involved in substrate binding, Hydrogen Deuterium Exchange (HDX) Mass Spectrometry was used to determine solvent accessibility of SteC_210-457_ with and without the FMNL1 peptide. Detected SteC peptides covered the entire protein (**Figure S1e**). Amino acids 388-396 of SteC within αF of the kinase domain C lobe showed the strongest differential, indicating that this region becomes protected from solvent in the presence of FMNL1 peptide (**Figure 1c**). We conclude that the minimal kinase domain required for *in vitro* phosphorylation spans amino acids 210-429 of SteC with regions of the minimal C lobe essential for catalysis and substrate binding.

### SteC kinase domain is monomeric in solution

Recently, it was suggested, based on a dimeric crystal structure of an SteC construct encompassing amino acids 202-375, that the kinase domain of SteC functions through an unusual dimeric arrangement in which one monomer supports the catalytic function of the other^8^. As our data reveal an essential role for additional regions of the C lobe, we used size exclusion chromatography coupled to multi-angle laser light scattering (SEC-MALLS) to determine the oligomeric state of both inactive and active SteC constructs in solution. Analysis of SteC_210-375_, SteC_210-429_ and SteC_210-457_ at multiple concentrations clearly demonstrated that each fragment is monomeric in solution (**Figure 1d**). Therefore, even though dimerisation can activate or inactivate certain kinases^17^, our in-solution analysis of the active kinase domain suggests that dimerisation does not represent a key regulatory feature of the activity of SteC.

### SteC is phosphorylated at S379

Upon the addition of ATP, SteC becomes phosphorylated *in vitro* (**Figure 1b** and Poh et al^9^). To identify putative sites of phosphorylation that might mediate activation of the kinase, SteC_1-457_ and SteC_1-457;K256H_ were expressed and purified from insect cells (**Figure S2a**). Analysis by mass spectrometry revealed that wild-type SteC was phosphorylated at S379 upon expression and purification from insect cells (**Figure 2a**) and additionally at S75 upon the addition of ATP to an *in vitro* kinase reaction (**Table S2**). The kinase mutant variant, SteC_1-457;K256H_, was also phosphorylated at S379 during expression and purification, with a similar proportion of phosphorylated/unphosphorylated peptides identified (**Figure 2b**). *In vitro* incubation with ATP only caused a minimal change in the proportion of identified peptides containing S379 phosphorylation for either WT or K256H SteC. Together, this suggests that phosphorylation at S379 of SteC was mediated by a eukaryotic kinase during insect cell expression and is not primarily an autophosphorylation event. To investigate S379 phosphorylation further, bacterially expressed SteC_210-457_ or SteC_210-429_ was analysed next. In these conditions, the ratio of phosphorylated/non-phosphorylated S379-containing peptides was lower, at approximately 0.4 (**Figure S2b**). Interestingly, the ratio of phosphorylated peptides was not increased upon the addition of ATP, yet, no phosphorylated S379 peptides were identified when kinase dead SteC_210-457;K256H_ was analysed (**Figure S2b**). This suggests that during expression of SteC in *E. coli*, autophosphorylation occurs, possibly in a semi-unfolded ‘prone-to-autophosphorylation’ state^18^, but not when the kinase domain is fully folded.

**Figure 2:**
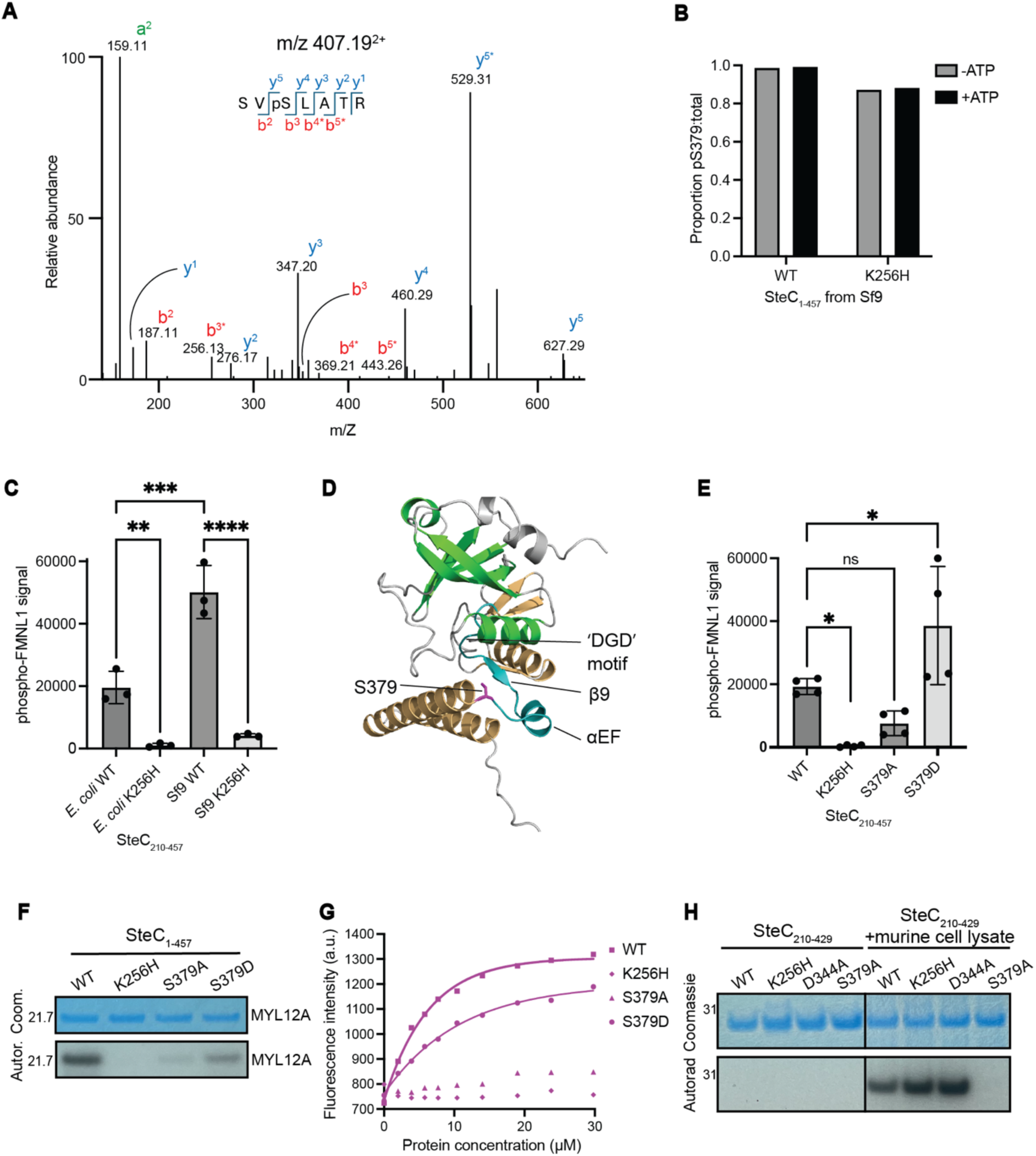
SteC is phosphorylated on S379 mediating activity and nucleotide binding. a) SteC_1-457_ expressed in Sf9 cells was analysed by mass spectrometry. Peptide SVSLATR (S377-R383) was detected in S379-phosphorylated and unphosphorylated forms following fragmentation obtained for precursor ion at m/z 407.2. C-terminal (y) ions and N-terminal (b) ions are presented here. y ions at 627 and 726 and the b3 ion at 354 indicate presence of phosphorylation. 529 represents unphosphorylated y5 minus H_2_O. b) S379 phosphorylated / non-phosphorylated peptides ratios were calculated as determined by mass spectrometry for the indicated conditions. c) Quantification of triplicate biological repeats for radioactive kinase assays performed with 5 μM SteC_210-457_ expressed in *E. coli* and Sf9 and 100 μM FMNL1 peptide (see also **Figure S2c**). Quantification was performed with ImageJ. d) AlphaFold2 prediction of the SteC kinase domain. N lobe secondary structural elements are highlighted in green and C lobe in light orange. The putative activation segment is highlighted in teal from D364 of the DGD motif to αF, including β9 and αEF. S379, at the C-terminal end of the activation segment, is shown as a purple stick. e) Quantification of four biological repeats of radioactive kinase assays containing 5 μM of SteC_1-457_ WT or associated mutants expressed in *E. coli* together with 100 μM FMNL1 peptide (see also **Figure S2d**). f) Radioactive kinase assays with SteC_1-457_ WT and mutants and MYL12A WT at 100 nM kinase and 5 μM MYL12A. Representative of 2 repeats. g) Fluorescence titrations of 500 nM mant-AMPPNP with SteC_210-429_ WT and mutants expressed in *E. coli*. The increase in fluorescence at increasing protein concentrations was measured. Data are representative of 2 repeats for each construct. Fitting curves are shown as purple lines. See also **Figure S2g**. h) Radioactive kinase assays of SteC_210-429_ WT and mutants expressed in *E. coli* at 5 μM with FMNL1 peptide at 100 μM or with 1 μL murine cell lysate (right). Statistical analysis used one-way ANOVA (*, p < 0.05; **, p < 0.01; ***, p < 0.001; ****, p < 0.0001; ns, non-significant) with Dunnett’s multiple comparisons test (C&E). Assumption of normally distributed data was accepted using a Shapiro-Wilk and Kolmogoro-Smirnov test.

### S379 phosphorylation is required for the activity of SteC

Analysis of the catalytic activity of SteC towards the FMNL1 peptide indicated that SteC_210-457_ expressed from insect cells was significantly more active than SteC_210-457_ expressed from bacteria (**Figure 2c** and **S2c**). S379 is located between αEF and αF of the C lobe, which encompasses regions of the kinase often referred to as the activation segment (**Figure 2d** and **S1b**). As phosphorylation of residues within the activation segment often control kinase function^19,20^, we hypothesised that phosphorylation of S379 might represent such a regulatory mechanism within SteC. To test whether phosphorylation of S379 activated kinase activity, mutation S379A, which cannot be phosphorylated and the phospho-mimetic variant, S379D, were introduced into SteC_210-457_ and proteins were expressed and purified from bacteria. As expected, WT SteC_210-457_ mediated phosphorylation of the FMNL1 peptide and SteC_210-457;K256H_ did not (**Figure 2e and S2d**). Phosphorylation of the peptide substrate was dependent on S379; becoming severely diminished upon incubation with SteC_210-457;S379A_, with phosphorylation not only restored, but increased when S379 was mutated to aspartic acid (**Figure 2e and S2d**). The increase in activity for the S379D variant of SteC was also evident upon analysis of SteC phosphorylation, which was barely detected for WT, catalytic mutant K256H or S379A variants, but evidently stronger in the S379D condition (**Figure S2d**). This finding revealed that additional, unidentified, SteC residue(s) beyond S379 and S75 become phosphorylated under *in vitro* conditions and led us to speculate that the requirement for S379 was not substrate specific. Indeed, whether analysing SteC_210-457_ or SteC_1-457_, *in vitro* phosphorylation of a second substrate, MYL12A^8^, was strongly dependent on S379 (**Figure 2f** and **S2e**). Therefore, we define S379 as a new phosphorylation site required for efficient activity of SteC towards known substrates.

### S379 phosphorylation mediates nucleotide binding

Next, we interrogated why S379 is important for the catalytic activity of SteC. Phosphorylation of the activation loop as a mechanism to activate protein kinases^19^ might occur through increased substrate binding^21^. To test whether S379 phosphorylation alters substrate binding we monitored the ability of the minimal active kinase domain, SteC_210-429_, to interact with an N-terminally biotinylated FMNL1 peptide by biolayer interferometry. We determined the following binding constants for SteC_210-429_ - WT K_d_ = 7.9 ± 1.4 μM, and variants K256H K_d_ = 22 ± 3 μM, S379A K_d_ = 21 ± 3 μM and S379D K_d_ = 14 ± 2 μM, respectively (**Figure S2f**). Given these similar affinities, we next tested the hypothesis that phosphorylation of S379 mediates allosteric conformational changes that alter nucleotide binding. This was monitored by analysing binding of a fluorescent non-hydrolysable ATP analog (mant-AMPPNP) to SteC_210-429_ by fluorescence spectroscopy. The data showed that WT SteC_210-429_ and SteC_210-429;S379D_ bound mant-AMPPNP with affinities of 5.4 ± 0.4 μM and 11.8 ± 1.5 μM respectively, whereas SteC_210-429;S379A_ showed significantly lower binding whereas SteC_210-429;K256H_ did not show any significant interaction (**Figures 2g and S2g**). From this, we conclude that phosphorylation of S379 activates SteC by increasing its affinity for ATP.

### S379 is phosphorylated by a mammalian kinase independent of SteC kinase activity

To explore the hypothesis that a mammalian kinase phosphorylates SteC, the phosphorylation of bacterially expressed SteC_210-429_ was analysed *in vitro* following incubation of recombinant protein with mammalian cell lysate. For this experiment an additional putative catalytic mutant, D344A, was analysed. Based on sequence and structural predications (**Figure 1a and S2b**), we hypothesised that D344 represents part of an HRD-like motif, which in the case of SteC, lacks the arginine, but retains the histidine and aspartate conserved in all eukaryotic protein kinases^14^. Phosphorylation of SteC_210-429_, SteC_210-429;K256H_ and SteC_210-429;D344A_ was observed but when S379 was mutated to alanine (SteC_210-429;S379A_) no phosphorylation was detected (**Figure 2h**). This demonstrates that a mammalian kinase can phosphorylate SteC at residue S379. Furthermore, it suggests that without S379 phosphorylation, any subsequent phosphorylation of SteC is not observed.

### S379 is required for SteC induced actin polymerisation in infected cells

SteC mediates the polymerisation of actin into dense foci associated with the *Salmonella* micro-colony^9^. To test whether S379 was required for dense actin foci formation during infection, 3T3 fibroblasts were infected with *steC* mutant bacteria expressing either HA-tagged WT or mutant SteC. Immunoblotting the pellet and the post-nuclear supernatant fractions of infected cells for HA and the *Salmonella* protein DnaK, demonstrated that each HA-tagged construct was expressed (pellet fraction containing bacteria) and translocated (supernatant fraction lacking bacteria) into the host cell cytosol (**Figure S3a**). As expected, cells infected with WT *Salmonella* showed association of actin, stained with phalloidin, at the microcolony (**Figure 3a**), and this occurred in approximately 40% of infected cells (**Figure 3b**). Actin polymerisation was significantly reduced when cells were infected with *steC* mutant *Salmonella* and restored upon expression of WT SteC from the bacteria (**Figure 3a,b**). Cells infected with *steC* mutant *Salmonella* strains expressing either SteC_K256H_ or SteC_S379A_ were unable to induce actin polymerisation. Expression of SteC_S379D_ from *steC* mutant bacteria restored the ability of SteC to induce dense actin foci around the micro-colony to levels that were similar to cells infected with bacteria expressing WT SteC (**Figure 3a,b**). We conclude that phosphorylation of S379 is essential for SteC-induced actin polymerisation during infection.

**Figure 3:**
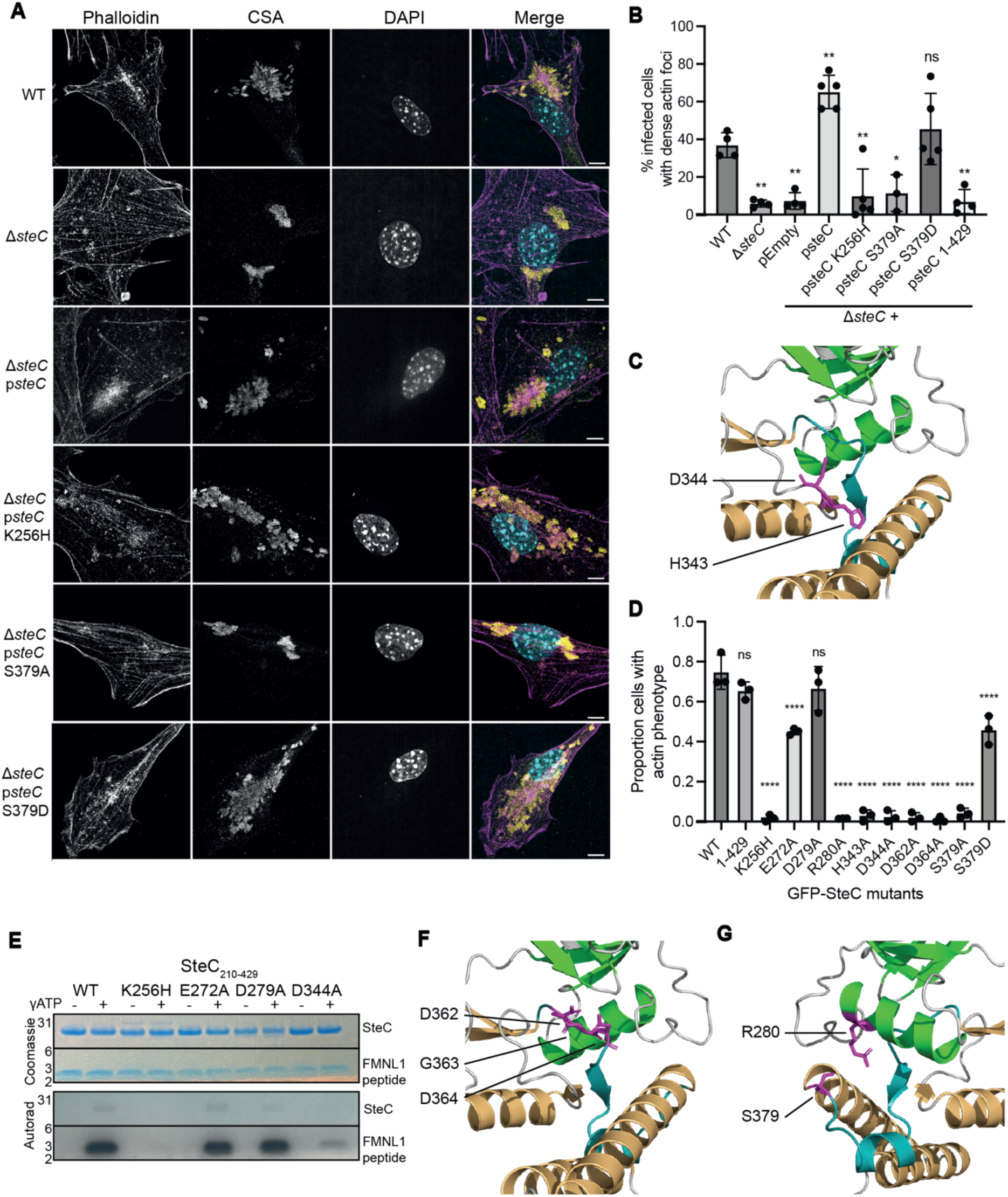
S379 is essential for SteC induced actin polymerisation during infection. a) Representative super-resolution images of Swiss 3T3 mouse fibroblasts infected with *ΔsteC Salmonella* strains expressing the indicated SteC variants. Scale bars represent 5 μm. In merge panel, phalloidin is in magenta, CSA (*Salmonella*) in yellow and DAPI in cyan (see also **Figure S3a**). b) For each *Salmonella* strains indicated 3-5 coverslips from two independent infections were prepared and 100 infected 3T3 cells were scored (blind) for the formation of dense actin polymerisation associated with the *Salmonella* microcolony. Each data point represents a coverslip and bars represent the mean and standard deviation. c) AlphaFold2 prediction of SteC kinase domain with the putative activation segment highlighted in teal from D364 including the DGD motif, β9, αEF and S379 prior to αF, N-lobe in green and C-lobe in sand. H343 and D344 are shown in purple sticks, indicating their position in relation to the catalytic left and activation segment. d) HEK 293ET cells were transfected with vectors expressing SteC WT and mutants. 100 transfected cells from each of 3 coverslips (representing biological repeats) were blind counted (see also **Figure S3b**). Each dot represents a coverslip and bars represent mean and standard deviation. e) Radioactive kinase assay of SteC_210-429_ WT and mutants expressed in *E. coli* at 5 µM with 100 µM FMNL1 peptide. Data representative of 2 repeats. f) Same as Figure 3c, but with D362, G363 and D364 shown in purple sticks. g) As for **Figures 3c and 3f** rotated 180° in the y axis, with S379 and R280 shown in purple sticks, indicating their proximity (4 Å apart). Statistical analysis used one-way ANOVA (*, p < 0.05; **, p < 0.01; ***, p < 0.001; ****, p < 0.0001; ns, non-significant) with Dunnett’s multiple comparisons test (B&D). Assumption of normally distributed data was accepted using a Shapiro-Wilk and Kolmogoro-Smirnov test.

### SteC contains non-canonical catalytic motifs in its depleted C lobe

So far, our data reveal a new phosphorylation site of SteC, S379, that mediates kinase activity. Kinases that require activation loop phosphorylation usually have an arginine residue adjacent to the catalytic aspartate and this forms the so called “HRD” motif within the C lobe of the kinase domain^14^. The aspartate is the most conserved residue of this motif and interacts directly with the substrate to orientate the hydroxyl acceptor group. Dai et al^8^ proposed D364 performs this function, yet this conclusion is based on the structure of an inactive form of SteC. As noted above, SteC contains an HD at residues 343-344 (**Figure S1b**) and we therefore hypothesised that this motif, positioned just after β6, might be required for function (**Figure 3c**). Cells expressing GFP-tagged SteC showed evident actin foci formation, and as expected this was absent in cells that expressed SteC_K256H_ or SteC_S379A_ (**Figure 3d and S3b**). Expression of either SteC_H343A_ or SteC_D344A_ was unable to induce actin foci formation, suggesting that these amino acids are required for the catalytic potential of SteC. Indeed, recombinant SteC_210-429_ with an alanine substitution at D344 showed minimal phosphorylation of the FMNL1 peptide, demonstrating the importance of this residue for SteC activity and function (**Figure 3e**).

Interestingly, in line with the observations by Dai et al.^8^, mutation of D362 or D364 to alanine also ablated SteC induced actin polymerisation whereas mutation of D279 did not (**Figure 3d and S3b**). This D_362_GD motif is positioned just after β8, where the canonical DFG motif, which mediates interaction with the Mg^2+^ ion to aid positioning of the gamma-phosphate, is found (**Figure 3f** and **Figure S1b**). This raises the intriguing possibility that D_362_GD might represent a non-canonical DFG motif.

Surprisingly, the previously predicted glutamic acid, E272^9^, which would normally stabilise the invariant lysine in subdomain II and ATP, was not required for the function of SteC (**Figure 3d and S3b**), nor its catalytic activity towards the FMNL1 peptide (**Figure 3e**). One possible explanation for this is that E272 is found outside alpha helix (αC) of subdomain III, where the αC glutamate would normally be positioned (**Figure S1b** and PDB: 8JBI^8^). Regardless, it remains unclear how SteC fulfils this important function.

Finally, we tested the hypothesis that R280, which has a positive charge and resides on αC helix (**Figure S1b**), might be required to stabilise the negative charge upon S379 phosphorylation (**Figure 3g**). GFP-tagged SteC_R280A_ was unable to induce actin polymerisation supporting this hypothesis (**Figure 3d and S3b**). We therefore propose that even though SteC lacks a classical HRD motif, the phosphate of pS379 might instead interact with R280 as a means of allosterically altering the active site to facilitate catalysis. Ultimately, why the putative ‘HD motif’, which includes the catalytically important D344, is required for function, and whether D362 and D364, which are also required for function, stabilise the divalent cation in a DFG-like role, requires further investigation.

### S379 is conserved across *Salmonella* species and bacterial SteC homologs

SteC is highly conserved in most *Salmonella* serovars (**Figure 4a**). Analysis of 879 clinical isolates of *Salmonella* revealed that K256, H343, D344, D362, D364 and S379 residues are all conserved (**Figure S4a**). Furthermore, aligning of putative SteC homologues from *Yokenella regensburgei* (98% coverage, 41% amino acid identity), *Cedecea neteri* (92%, 42%), *Sodalis praecaptivus* (86%, 34%) and *Erwinia mallotivora* (42%, 49%) showed that these six residues are conserved across all the homologues (**Figure S4b**). These findings suggest a common catalytic and phosphorylation-regulated mechanism of SteC across diverse bacterial species.

**Figure 4:**
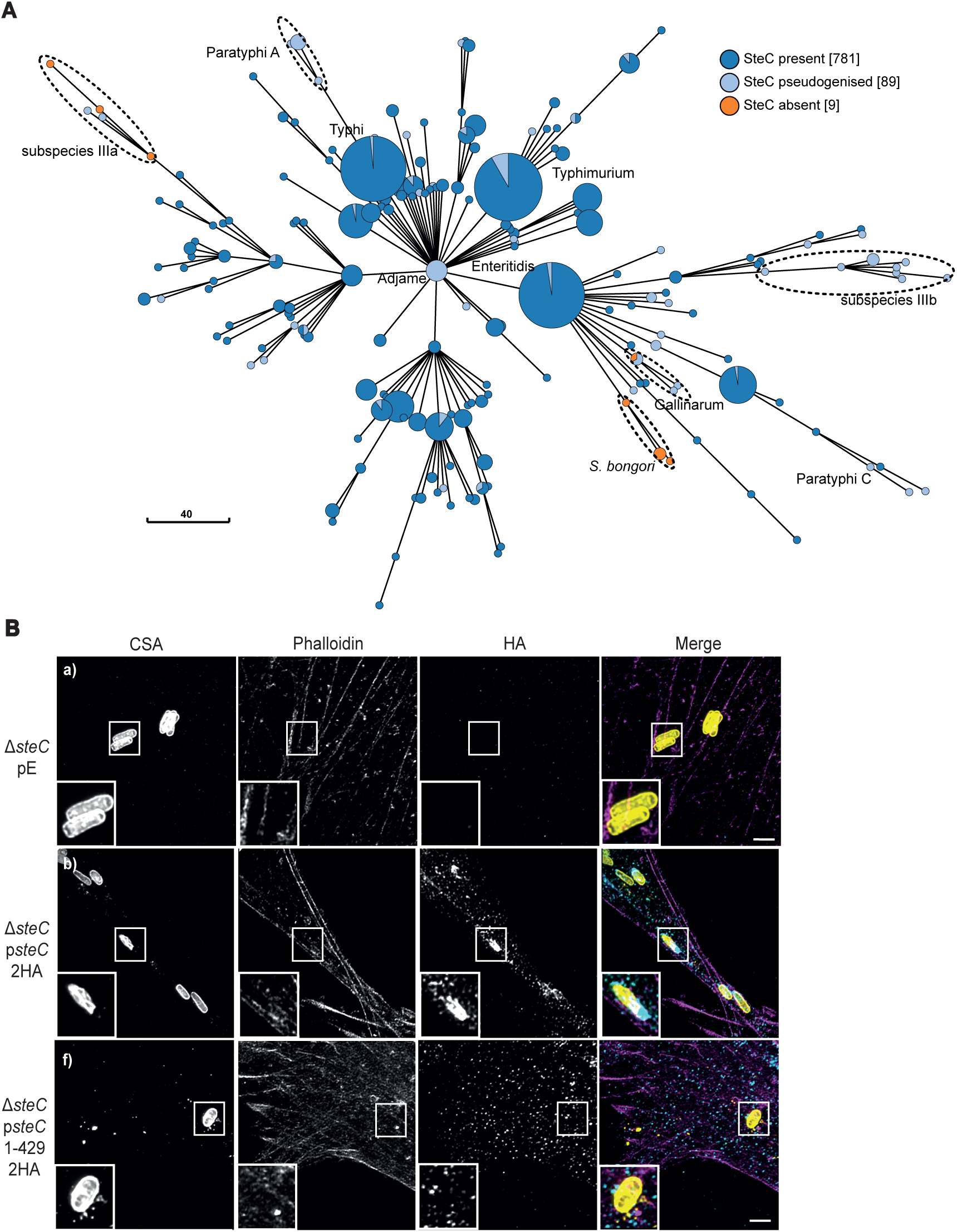
S379 and key catalytic motifs are conserved in SteC and its bacterial homologs. a) GrapeTree^25^ visualisation of SteC distribution in 879 *Salmonella* complete genomes. The MStree was constructed using the rMLST scheme, which could differentiate *Salmonella* at serovar level based on ribosome loci. Each bubble represents a distinct rST. The branch lengths correspond to the number of allele differences at the ribosome loci. Nodes with only a single allele difference were collapsed into bubbles, which exhibit high concordance with serovars. The size of each bubble is proportional to the number of genomes it represents. The colours of the bubbles represent presence, absence or pseudogenisation of SteC. The phylogenetic analysis is not rooted, meaning evolutionary relationships cannot be inferred. b) Representative super resolution microscopy analysing subcellular localisation of HA tagged SteC variants following the infection of 3T3 cells. CSA labels *Salmonella* and Phalloidin stains actin. In the merge, CSA is shown in yellow, phalloidin in magenta and HA in cyan. See also **Figure S5**. Scale bars represent 5 μm.

Interestingly, however, the C-tail was not highly conserved in the putative homologues (**Figure S4b**). We noted that unlike WT SteC, translocation of SteC_1-429_, which lacks the C-tail, was unable to polymerise actin (**Figure 3b and S3a**). Yet, when assayed *in vitro*, SteC_1-429_ phosphorylates the FMNL1 peptide (**Figure 1b**) and when ectopically expressed in HEK 293ET cells it also retains activity (**Figure 3d and S3b**). To investigate this further, the subcellular localisation of SteC was analysed using super resolution microscopy. This revealed that WT SteC formed structures that colocalised with actin foci or the bacteria. In contrast, in 3T3 cells infected with bacteria expressing SteC_1-429_, SteC_1-429_ was localised in distinct puncta across the cell (**Figure 4b and S5**). We therefore hypothesise that during infection, the C-tail of SteC is an essential regulator of SteC function, possibly by contributing to its correct subcellular localisation.

## CONCLUSION

Here we reveal that instead of being activated through dimerisation or interaction with a binding partner, as suggested by previous models and observed for the minimal kinase *Shigella* effector OspG^15^, the kinase activity of SteC is induced through phosphorylation by a host kinase. Phosphorylation of SteC at S379, which is essential for actin polymerisation, represents a catalytically activating event that induces a dramatic increase in kinase domain nucleotide binding affinity. Rather than through a canonical HRD motif, this phosphorylation-dependent activation might instead be mediated through interaction with R280, present on the αC helix. Determining whether this stabilizes the hinge between the N and C lobes or triggers allosteric changes that enhance activity will likely require a full structural analysis of the kinase module bound to a substrate. However, to date, protein stability has represented a limiting step towards achieving this. Furthermore, whether the N-terminal portion of SteC (amino acids 1-193) contributes to kinase activity or substrate selectivity remains enigmatic.

Despite these remaining questions, this study revealed how phosphorylation of S379 represents an unappreciated mechanism of activating the only eukaryotic-like kinase from *Salmonella*. Several *Salmonella* effectors are controlled by host-mediated PTMs. SifA, for example is prenylated and S-acetylated by host machinery which leads to both increased membrane binding and *Salmonella* survival in infected mice^22^, and is also cleaved by caspase 3 to be activated^23^ The SseI effector undergoes S-palmitoylation^24^, which regulates its plasma membrane localisation and function. It is intriguing that the C-tail of SteC, which is not required for its kinase activity *in vitro*, is essential for actin polymerisation in infected cells, possibly by directing the correct subcellular localisation. Altogether, the reliance of modification(s) by host enzymes might represent a mechanism to ensure that the activity of a given effector is only unleashed in the host and only in the correct subcellular localisation within the host.

## Resource availability

Requests for further information and resources should be directed to and will be fulfilled by the lead contact, Katrin Rittinger. Stable reagents generated in this study will be made available on request. The HDX mass spectrometry proteomics data have been deposited to the ProteomeXchange Consortium via the PRIDE partner repository with the dataset identifier PXD061217. All other data reported in this study is available in associated source data files or will be shared by the lead contact upon request.

## Acknowledgements

We thank Ian Taylor for help with SEC-MALLS experiments, Dhira Joshi from the Francis Crick Institute Chemical Biology STP for peptide synthesis, Steve Howell, Tania Auchynnikava and Mark Skehel from the Francis Crick Institute Proteomics STP for mass spectrometry analysis; Matt Renshaw and Donald Bell from the Francis Crick Institute Light Microscopy STP for expert technical support; Yizhou Huang, Paul O’Sullivan, Magdalena Szczesna and Ioanna Panagi for advice on methods and Peter Hill for critical reading of the manuscript.

This work was funded by the Francis Crick Institute, which receives its core funding from Cancer Research United Kingdom (CC 2075), the United Kingdom Medical Research Council (CC 2075), and the Wellcome Trust (CC 2075); a Biotechnology and Biological Sciences Research Council (BBSRC) David Phillips Fellowship (BB/R011834/1) awarded to TLMT; a Medical Research Council (MRC) grant MR/V031058/1 awarded to TLMT and KR, which also funded IDDO; an Engineering and Physical Sciences Research Council (EPSRC) grant EP/X02377X/1, underwriting European Research Council Starting Grant, Re-kin awarded to TLMT, which also supported ML and a Wellcome Trust Investigator award to JCDH (Grant number 222528/Z/21/Z).

For open access, the author has applied a CC BY public copyright license to any author-accepted manuscript version arising from this submission.

## Author Information

Conceptualization: TLMT and KR; Methodology: TLMT, TDP, JCDH and KR; Investigation and data analysis: TDP, BL, DE, IDDO, JH, ML, SM, LM, YL and XY; Writing – Original Draft: TDP, TLMT and KR; Writing – Review & Editing: all authors; Funding Acquisition: KR and TLMT; Supervision: XY, JCDH, TLMT and KR.

## Supplemental information

Document S1. Figure S1-S5 and Tables S1-2

## STAR Methods

### Key resources table

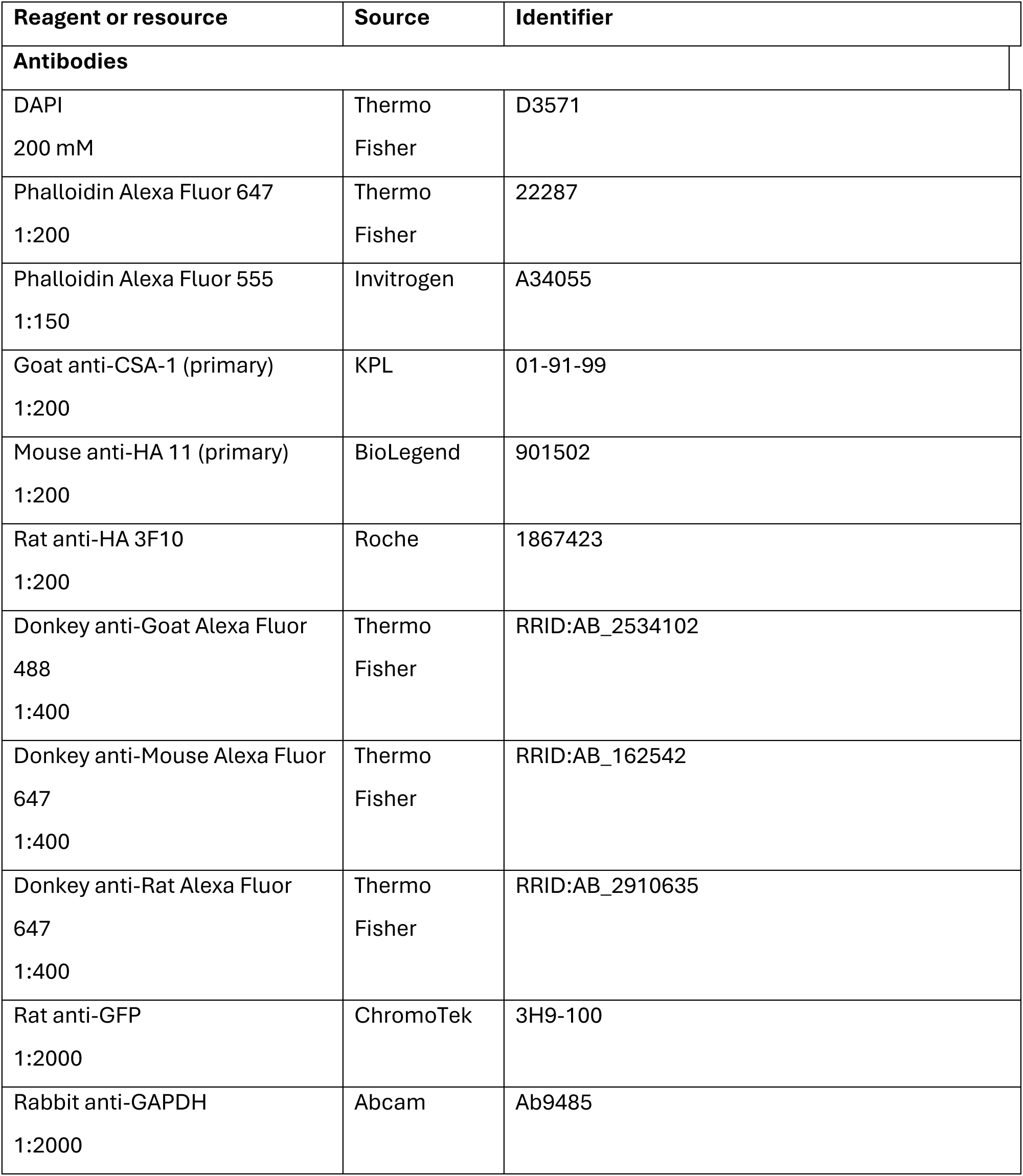
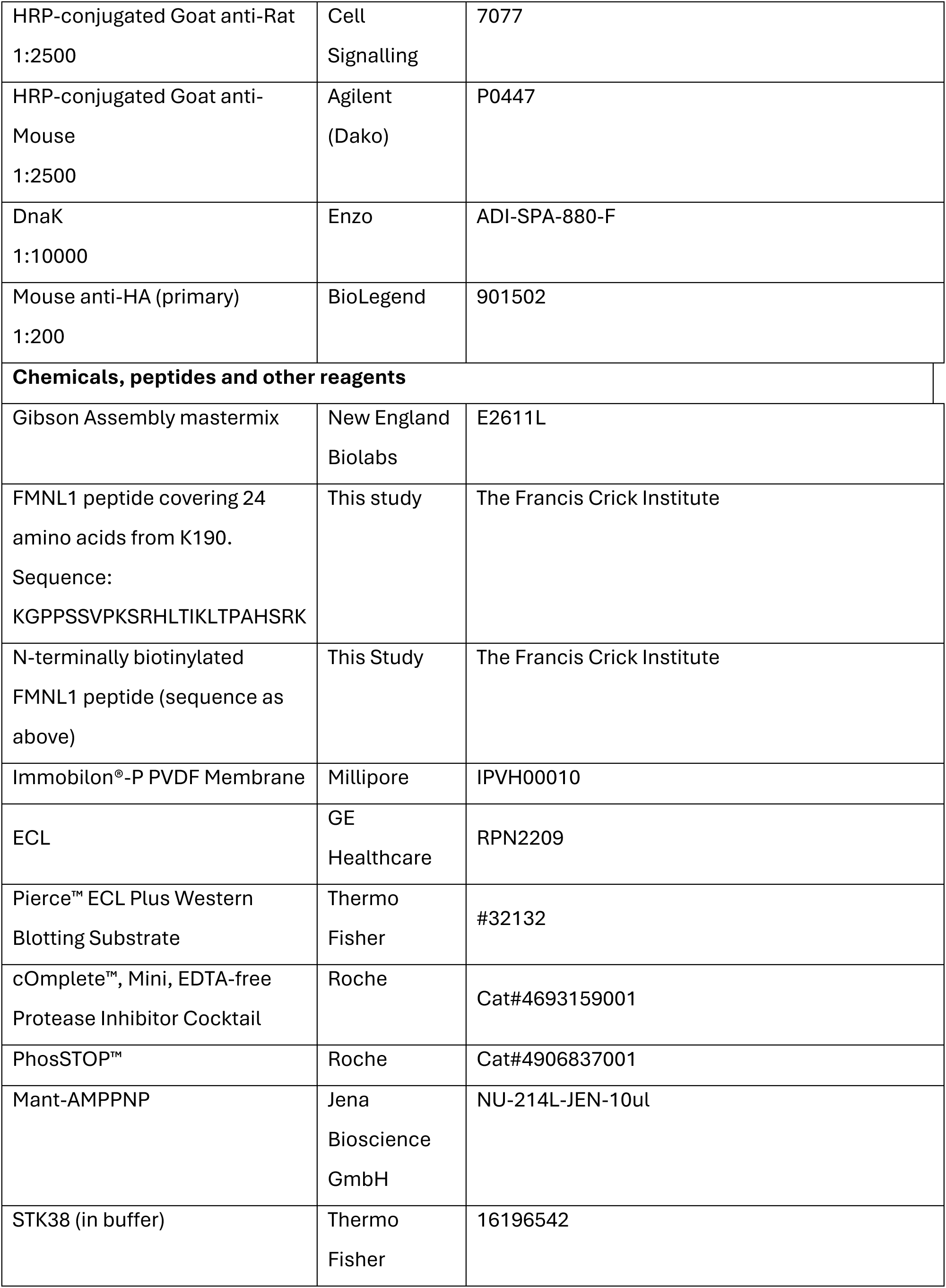
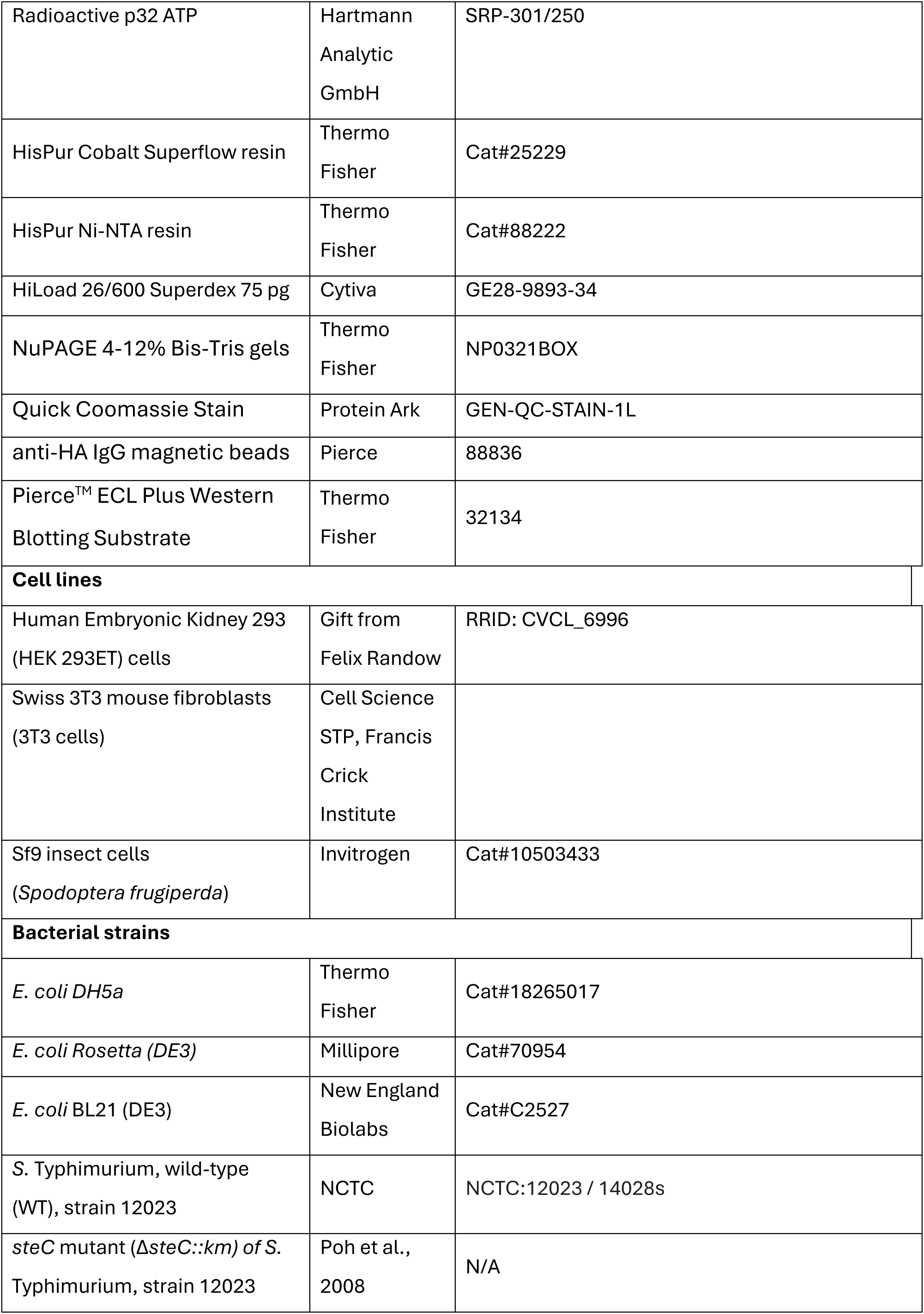
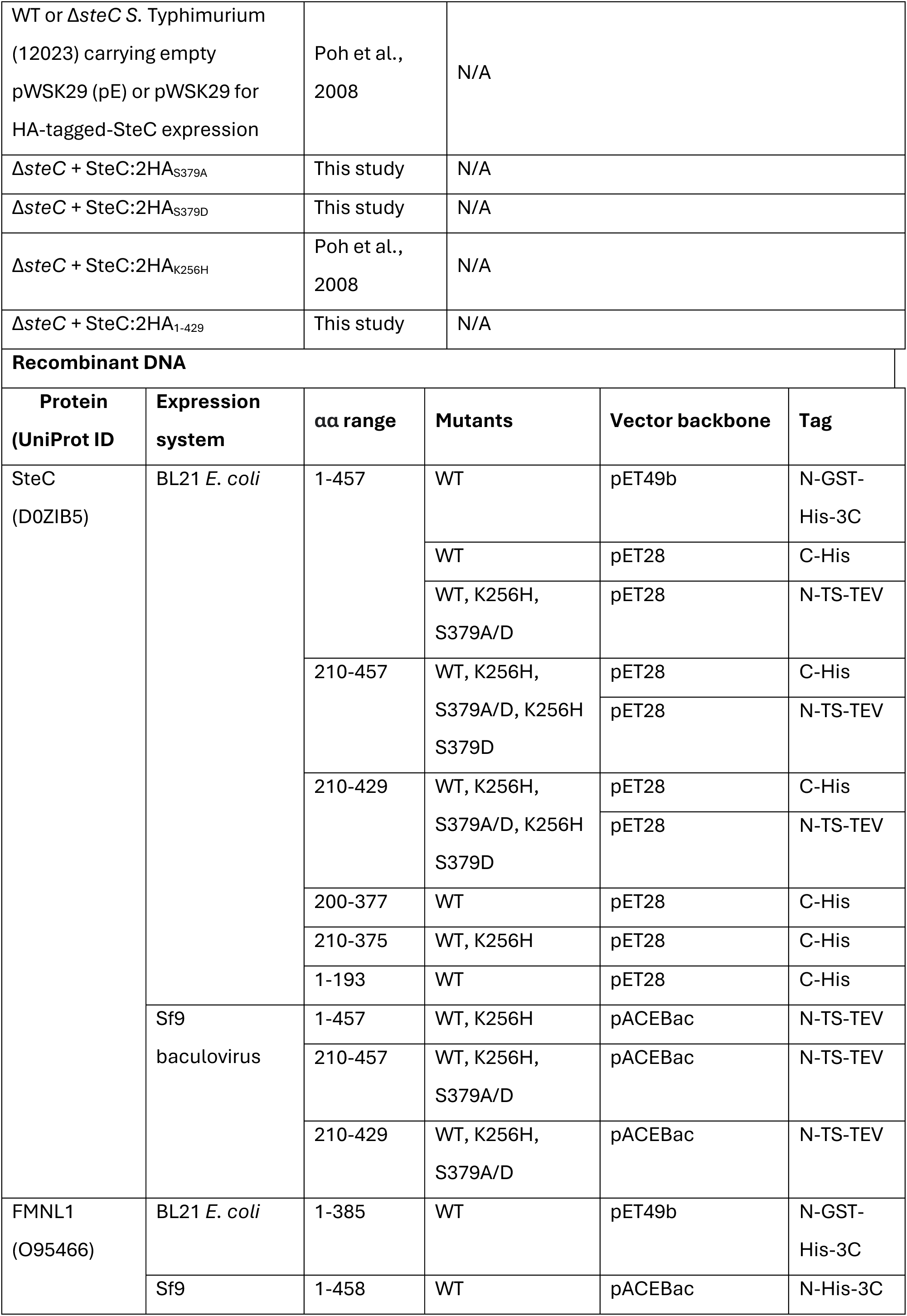
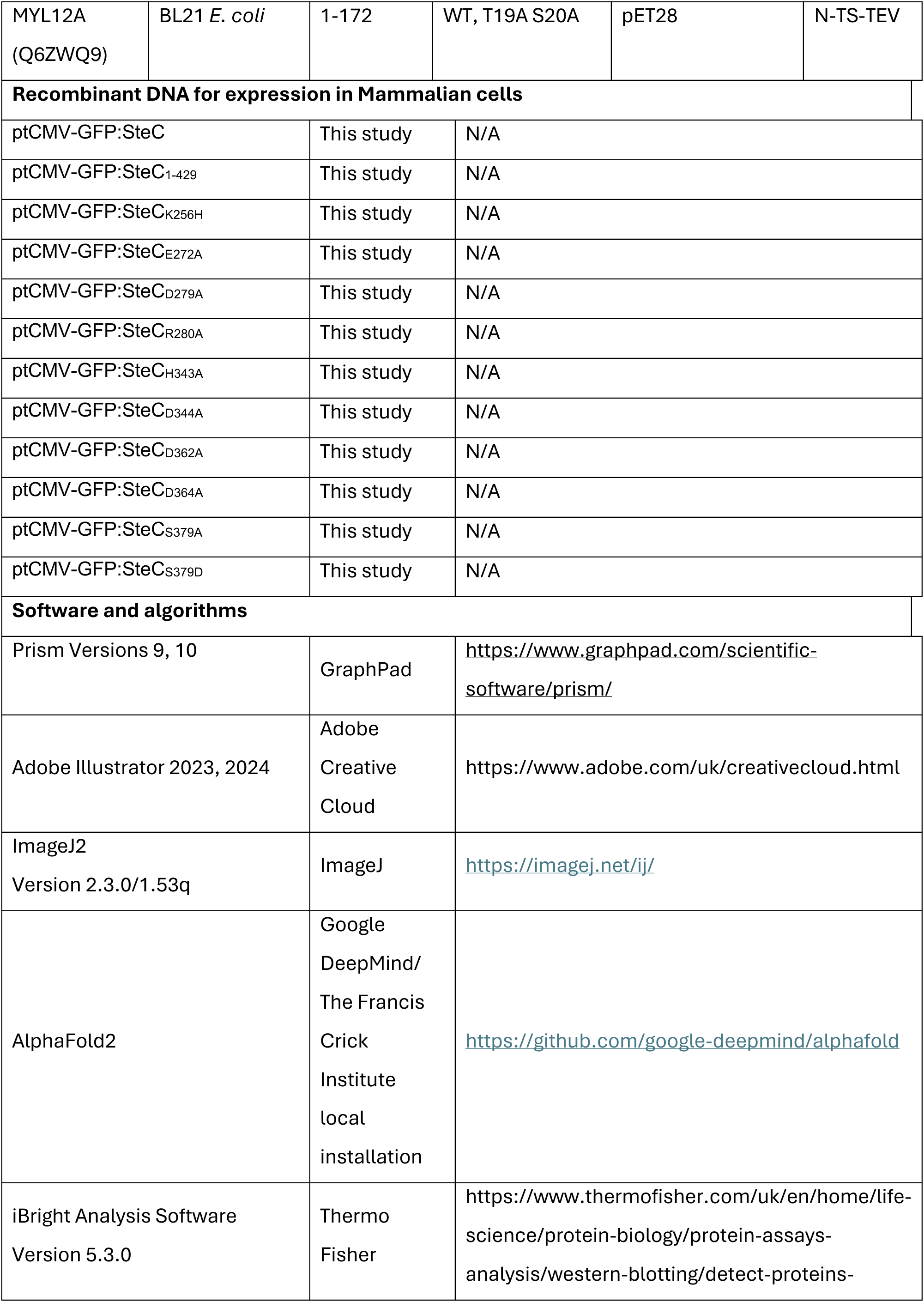
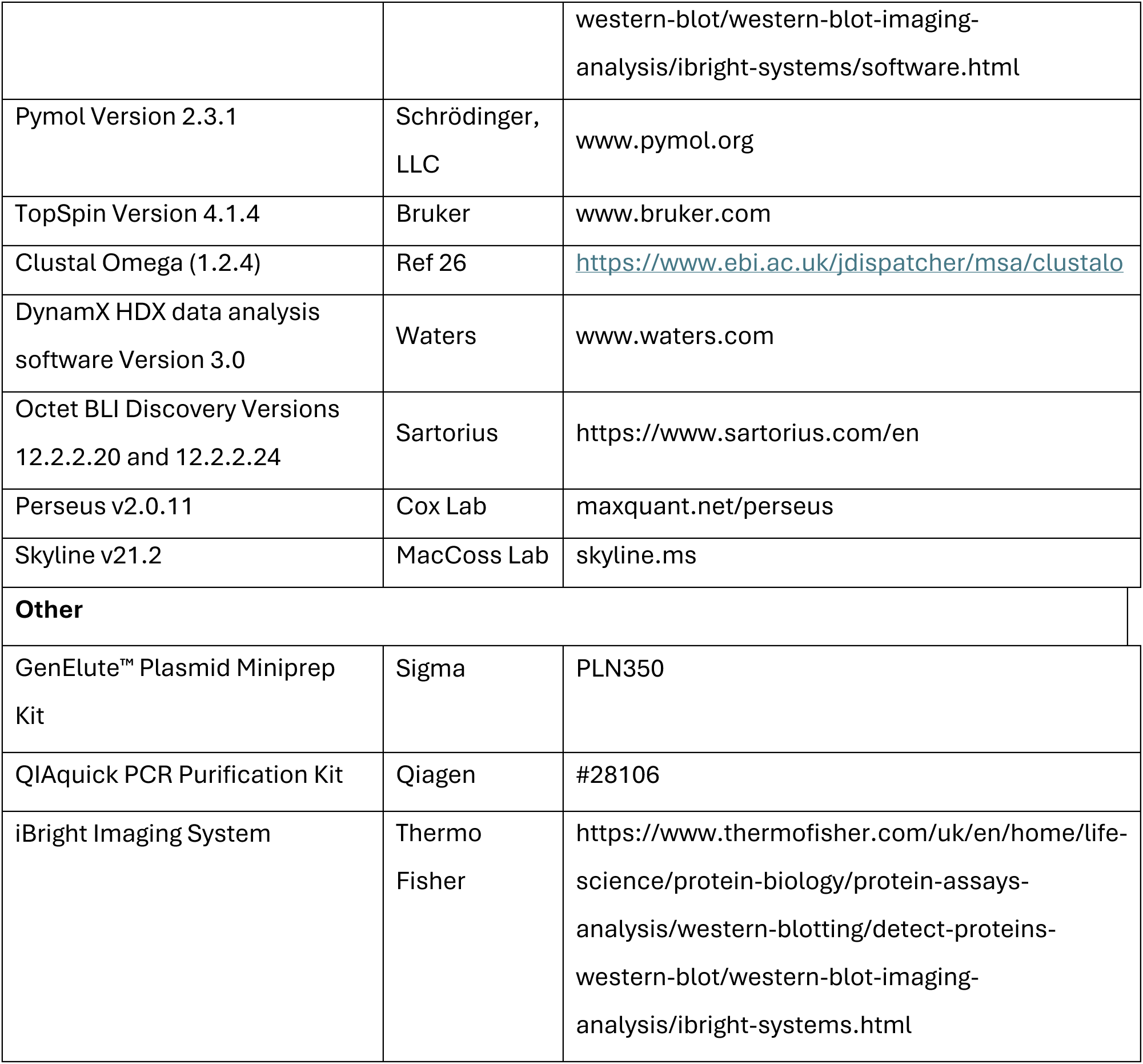

### Methods

#### Protein structure prediction

Experimentally derived protein structures were retrieved from the Protein DataBank (PDB) (https://www.rcsb.org/). AlphaFold2 (AF) predicted structures were downloaded from the AlphaFold Protein Structure Database (https://alphafold.ebi.ac.uk/). The *steC* gene sequence with UniProt reference D0ZIB5 from *Salmonella* Typhimurium strain 14028s was used for all bioinformatic analyses and protein expression and purification. PDB and predicted structures were visualised using Pymol (https://www.pymol.org/).

#### Plasmid design and cloning

Expression plasmids were produced using Gibson assembly^26^ (key resources table). Vector backbones used were pET28 (either with C-terminal uncleavable His6 or N-terminal Twin-Strep tag with TEV protease cleavage site for *E. coli* expression), pACEBac (either N-terminal His6-3c cleavage site or N-terminal Twin-Strep TEV protease cleavage site for insect cell expression), pTCMV (N-terminal GFP tag, for ectopic expression in mammalian cells) and pWSK29 (C-terminal 2HA tag for transformation of *Salmonella*). For expression of SteC and associated variants (K256H, S379A, S379D, and SteC_1-429_) from *Salmonella*, the *steC* open reading frame and C-terminal 2HA tag, was synthesised as a G-block from IDT technologies and inserted by Gibson Assembly into pWSK29 plasmid, which already contained the SteC promoter. Where required, mutagenesis was performed using the Quickchange protocol.

#### Protein expression and purification

Competent BL21 (DE3) *E. coli* cells were transformed with relevant plasmids, grown to mid-log phase at 37°C in LB, measured by OD_600_ of 0.7-1, and expression was induced with 0.5 mM Isopropyl-β-D-thiogalactoside (IPTG) at 20 °C. After 16-24 hours of expression, cells were pelleted and either frozen for future use or lysed directly. To lyse, the fresh or frozen cells were resuspended in a buffer of 50 mM HEPES (4-(2-hydroxyethyl)-1-piperazineethanesulfonic acid) pH 7.5, 300 mM NaCl, 10% glycerol (v/v), 10 mM MgCl_2_, 20 mM imidazole (for His-tagged proteins only), 0.5 mM tris(2-carboxyethyl)phosphine hydrochloride (TCEP), 1 x protease inhibitor cocktail (Roche) and 2 μg/ml DNAse (Merck). Lysis was performed by sonication. The lysate was cleared by ultracentrifugation at 48,000 x g for 1 hour at 4 °C. Lysate supernatant was passed over affinity resin, either using nickel nitrilotriacetic acid (NiNTA) or Streptactin resin affinity chromatography, in a gravity column at 4 °C. Resin washing was performed with Buffer A (50 mM HEPES pH 7.5, 300 mM NaCl, 10% glycerol plus 20 mM imidazole for His-tagged proteins) and high salt (50 mM HEPES pH 7.5, 1 M NaCl) prior to elution with buffer B (50 mM HEPES pH 7.5, 300 mM NaCl, 10 mM MgCl_2_, 10% glycerol, 0.5 mM TCEP and either 300 mM imidazole or 2.5 mM desthiobiotin). All proteins were purified at 4°C and 10% glycerol added to prevent aggregation.

For insect cell expression, EmBacY cells were transformed with the pACEBac plasmid by electroporation pulsed at 2 kV with a Gene Pulser II (BioRad) and incubated at 37 °C for 6 hr to stimulate transposition. 100 μL of these cells were incubated for 48 hr on plates with antibiotics selecting for the bacmid, the transposed plasmid and a helper plasmid. DNA was extracted from positive colonies and transfected into Sf9 cells. Cells expressing GFP denoted the production of viable virus. Virus was amplified to reach the potency required for adequate protein expression. The P2 or P3 virus was deemed potent when > 80 % of cells were infected and the cell viability was 85-92 %. Sf9 at 2 x 10^6^ cells/ml were infected with potent virus at a ratio of 500:1 and incubated for 72 hours at 28 °C with shaking. Protein derived from insect cells was purified as described for *E. coli* expressed protein with the following small adjustments: Lysis buffer included 0.1 mM 4- (2-aminoethyl)benzenesulfonyl fluoride hydrochloride (AEBSF) serine protease inhibitor. Cell lysis was performed with 0.1% Triton added to the lysis buffer with stirring at 4 °C for 45 min. For resin affinity purification, batch washing by resin centrifugation was performed prior to gravity column washing due to the viscosity of the lysate. Finally, elution of His-tagged proteins expressed in Sf9 required serial elution with buffers containing 300 mM and 500 mM imidazole.

For SteC kinase domains and MYL12A, size exclusion chromatography (SEC) was the final purification step. SEC buffer comprised 50 mM HEPES pH 7.5, 150 mM NaCl, 10 mM MgCl_2_, 0.5 mM TCEP and 10% glycerol (v/v), unless otherwise stated. For large scale protein purifications, HiLoad 16/600 Superdex (200 or 75) columns were used on an AKTA prime system at 4 °C with a flow rate of 1 ml/min with 2 ml fractionation. For smaller scale purifications, Superdex 200 or 75 Increase 10/300 GL SEC columns were used, with a flow rate of 0.5 ml/min with 500 μL fractionation. In the cases of SteC full length and FMNL1_1-458_, SEC was not possible due to protein aggregation and proteins were dialysed into SEC buffer overnight at 4 °C. Protein was concentrated using Amicon Ultra centrifugal filters, of 50, 30 or 10 kDa filter sizes. Protein concentration was measured using absorbance at 280 nm on a Nanodrop instrument. Aliquots of protein were flash frozen in liquid nitrogen and stored at −80 °C. For biophysical techniques potential aggregation was cleared by ultracentrifugation at 17,000 x g at 4 °C for 10 min, protein dialysed where necessary, and the concentration re-estimated using a Jasco V-760 Spectrophotometer.

#### 1D NMR

One-dimensional ^1^H nuclear magnetic resonance (1D NMR) protein spectra were recorded at 25 °C on Bruker AVANCE spectrometers operating at 800 MHz in NMR buffer (50 mM HEPES pH 7.5, 150 mM NaCl, 10 mM MgCl2, 0.5 mM TCEP and 5% D_2_O). Data were acquired and processed with Topspin (Bruker).

#### Hydrogen-deuterium exchange mass spectrometry

Hydrogen-deuterium exchange mass spectrometry (HDX-MS) was performed with 5 µL of 5 µM proteins (individually or in combination) incubated with 40 µL of D_2_O buffer at room temperature for 3, 30, 300 and 3000 seconds in triplicate. The labelling reaction was quenched by adding chilled 2.4% v/v formic acid in 2 M guanidinium hydrochloride and immediately frozen in liquid nitrogen. Samples were stored at −80 °C prior to analysis. The quenched protein samples were rapidly thawed and subjected to proteolytic cleavage by pepsin followed by reversed phase high-performance liquid chromatography (HPLC) separation. Briefly, the protein was passed through an Enzymate ethylene-bridged hybrid (BEH) immobilised pepsin column (Waters, UK) at 200 µL/min for 2 min and the peptides trapped and desalted on a 2.1 x 5 mm C18 trap column (Acquity BEH C18 Van-guard pre-column, 1.7 µm, Waters, UK) then eluted. Peptides were separated on a reverse phase column (Acquity UPLC BEH C18 column 1.7 µm, 100 mm x 1 mm (Waters, UK). Peptides were detected on a Cyclic mass spectrometer (Waters, UK). Peptide identification was performed by MS^e 27^. The resulting MS^e^ data were analysed using Protein Lynx Global Server software (Waters, UK). Mass analysis of the peptide centroids was performed using DynamX software (Waters, UK). All time points in this study were prepared at the same time and individual time points were acquired on the mass spectrometer on the same day.

#### Size exclusion chromatography coupled to multi-angle laser light scattering

Size exclusion chromatography coupled to multi-angle laser light scattering (SEC-MALLS) was performed with 100 μL protein samples at concentrations of 0.25, 0.5, 1, 2, 4 and 8 mg/ml were first applied to a Superdex 200 10/300 INCREASE GL column equilibrated in 50 mM HEPES pH 7.5, 150 mM NaCl, 10 mM MgCl_2_, 0.5 mM TCEP and 3 mM NaN_3_ to separate species by hydrodynamic volume. Scattered light intensity was measured using a DAWN HELEOS II laser photometer, and dRI was measured using an OPTILAV-TrEX differential refractometer. The weight-averaged molecular mass of proteins was determined using the ASTRA software version 7.0 (Wyatt Technology Corp., Santa Barbara, CA) assuming Dn/dc to be 0.186 mL/g.

#### Radioactive kinase assays

Reactions were performed in standard SEC buffer. Each 15 μL reaction mixture contained 50 nM - 10 μM kinase (details stated in Figure legends), 100 μM ATP, 20 kBq [γ-32P]ATP (Hartmann Analytic, 9.25 MBq, 25 μL pot, 5000 Ci/mmol). Given the aggregation-prone nature of SteC_210-457_, for concentrations below that detectable by Coomassie staining, protein aliquots were thawed, thoroughly mixed and concentration recalculated using Nanodrop measurement, prior to serial dilution to the concentration required. FMNL1 K190 24mer peptide (pep_K190) was used at 100 μM. Reactions were incubated at 30 °C for 30 min prior to SDS-PAGE analysis, Coomassie staining, gel drying and exposure to radiography film in the dark for between 30 min and 18 hours before developing. 1 μl Swiss 3T3 mouse fibroblast lysate (see below details of its preparation) was used to spike relevant kinase assay samples. Quantification of kinase assays was performed using band quantification in ImageJ.

#### Phosphorylation mass spectrometry

SteC_1-457_ WT and K256H and FMNL1_1-458_ expressed in insect cells were reduced and alkylated in-gel (10 mM TCEP, 40 mM chloroacetamide for 20 min at 70°C) prior to trypsin digestion (modified sequencing grade, Promega) in 10 mM NH_4_HCO_3_ overnight at 37°C. Acidified supernatant was separated by high-performance liquid chromatography and loaded into a Lumos Tribrid Orbitrap mass spectrometer (all Thermo Scientific). Raw files were searched using Maxquant (maxquant.org) against FASTA sequences of relevant recombinant constructs, recent downloads of UniProt baculovirus related databases and a common contaminants database and modified residues were assigned. Visualisation was in Perseus (maxquant.net/perseus) and Skyline (skyline.ms).

#### Biolayer interferometry

Bio-Layer Interferometry (BLI) was performed on an Octet Red instrument (Fortebio/Sartorius) operating at 25 °C. The N-terminally biotinylated FMNL1 peptide was synthesised by the Chemical Biology STP (Francis Crick Institute). The peptide was solubilised in, and proteins were dialysed into 50 mM HEPES pH 7.5, 150 mM NaCl, 10 mM MgCl_2_ and 0.5 mM TCEP. 0.05% Tween-20 was added to the samples prior to the experiments. Octet Streptavidin (SA) biosensors were loaded with the biotinylated peptide (1 μg/ml) and then exposed to SteC protein concentrations ranging from 7.5 nM to 200 µM. Association and dissociation curves were recorded for each concentration. Control experiments with no peptide loaded on the sensors were recorded to correct for non-specific interactions of SteC proteins with SA sensors. Data were analysed using Octet BLI Analysis software (Sartorius) and in-house software^28^. The equilibrium dissociation constant (K_d_) was determined from the instrument response against SteC proteins concentration using least squares non-linear regression. The reported error is the mathematical error of the fit.

#### Nucleotide binding

SteC proteins were analysed in SEC buffer. Mant-Adenylyl imidodiphosphate (AMPPNP) was acquired from Jena Bioscience GmbH. Data were collected on a Jasco FP-8500 Spectrofluorometer using an excitation wavelength of 355 nm and recording emission at 390-550 nm, in a 0.3 cm path length quartz cuvette (Hellma Analytics). Protein titrations were performed by recording full spectra and adding small volumes of protein to mant-AMPPNP at 500 nM. The signal at 442 nm was baseline subtracted and corrected for protein samples contributions and dilution, prior to plotting against the protein concentrations. Data were fitted using non-linear least squares regression with in-house software^28^. Average values and standard deviations were calculated from two independent measurements.

#### *Salmonella* strains

*Salmonella* WT 14028s and the same strain in which genomic *steC* had been replaced with a kanamycin cassettes were from Poh et al., 2008^9^. To generate *Salmonella* strains expressing SteC:2HA and associated variants from pWSK29 the *ΔsteC* strain was transformed by electroporation. *Salmonella ΔsteC* was grown in Luria Broth media (LB) to an OD_600_ of 0.4. The culture was cooled on ice for 30 min prior to washing the bacteria three times with autoclaved MilliQ-purified H_2_O and then autoclaved MilliQ H_2_O with 10% glycerol (once). The bacterial cell pellet was resuspended in 200 µl 10% glycerol and electroporation was performed with 50 µl of cell suspension and 100 ng DNA in a 0.2 mm cuvette at 2.5 kV, 200 ohms and 25 µF with a Pulse Controller Plus (BioRad). The product was incubated for 1 hr at 37 °C in 500 µl SOC and plated on LB agar plates with kanamycin and carbenicillin. Single colonies were selected and stored in 33% glycerol at −80 °C.

#### Cell lines and cell lysate preparation

Human Embryonic Kidney 293 cells (HEK 293ET cells) and Swiss 3T3 mouse fibroblasts (3T3 cells) were grown in Dulbecco’s Modified Eagle’s Medium (DMEM) (Sigma, USA) supplemented with 10% foetal bovine serum (FCS) (Gibco Life Sciences, UK) at 37 °C in 5% CO_2_. To produce murine cell lysate, two confluent 10 cm plates of 3T3 cells were washed in phosphate buffered saline (PBS) once then lysed in the plate with 1 ml Lumier ++ (Tris pH 7.4, 150 mM NaCl, 0.1% Triton, 1% EDTA, protease inhibitor cOmplete (Roche) and phosphatase inhibitor phoSTOP (Roche)) with 0.3% Triton for 15 min on ice, before pelleting at 17,700 x g for 10 min at 4 °C. 20 µl aliquots of the supernatant were stored at −20 °C.

#### *Salmonella* infection of mammalian cells

Overnight cultures of *S.* Typhimurium strains were diluted 1:33 in fresh LB and grown for 3.5 hrs at 37 °C with shaking (200 rpm) to obtain logarithmic phase bacteria. Cultures were added directly to 3T3 cells at a multiplicity of infection of 100:1. Bacterial invasion was allowed to proceed for 20 min at 37 °C in 5% CO_2_ after which the cells were washed twice in PBS and fresh media containing 100 μg / ml gentamicin was added for one hour prior to exchange with media containing 20 μg / ml gentamicin for the remainder of the experiment.

#### *In cellulo* analysis of SteC translocation

3T3 cells seeded on 6-well plates were infected with *Salmonella* WT, or the indicated *ΔsteC* strains expressing HA-tagged variants of SteC. Eight hours after invasion, cells were washed in PBS and lysed with 50 μl Lumier ++ (Tris pH 7.4, 150 mM NaCl, 0.1% Triton, 1% EDTA, protease inhibitor and phosphatase inhibitor) for 10 min on ice and centrifuged at 17,700 x g at 4 °C. After centrifugation the soluble layer representing the post-nuclear supernatant (PNS) was denatured by addition of 20 μl of 5x SDS buffer (125mM Tris-Cl pH 6.8, 4 % SDS, 10% glycerol, bromophenol blue and 5% β-mercaptoethanol). The pellet fraction, containing bacteria and mammalian cell nuclei, was denatured with addition of 100 μl of 2x SDS loading buffer. Samples were then boiled at 95°C for 7 min. Pellet samples were sonicated briefly to reduce viscosity prior to SDS-PAGE and immunoblot analysis with antibodies against HA, DnaK and GAPDH (Key resources table).

#### Immunoblotting

Denatured protein samples were run on 10-14% polyacrylamide gels by electrophoresis. A constant voltage of 110 V was applied for 90 min. Proteins were transferred onto a 0.2 μm polyvinylidene fluoride or polyvinylidene difluoride (PVDF) transfer membrane (Millipore) using a Trans-Blot® Turbo^TM^ Transfer System (BioRad). Membranes were blocked in 5 ml 5% bovine serum albumin (BSA) in Tris-buffered saline with 100 mM Tris pH 7.4, 150 mM NaCl, 0.1% Tween 20 (TBS-T) for 30 min at room temperature on a roller. The membranes were incubated with primary antibodies in 5% BSA overnight at 4 °C followed by three washes in TBS-T before being exposed to the appropriate secondary antibody in 5% BSA for 1 hr at room temperature. Next, after three further washes in TBS-T, membranes were incubated with either ECL^TM^ detection reagents or Pierce^TM^ ECL Plus Western Blotting Substrate and imaged on an iBright FL1500 Imaging System (Invitrogen, USA). See antibody list for details (key resources table).

#### Transfection of mammalian cells

HEK 293ET cells were seeded onto sterile glass coverslips coated with Poly-L-lysine (Sigma, USA) 24 hours prior to transfection. The cells were transfected with 3 μL Lipofectamine 2000 (Invitrogen, USA) and 600 ng of the indicated pTCMV GFP-SteC mutant vectors (key resources table) in 250 μL Opti-MEM Reduced Serum Medium (Gibco, UK). The cells were incubated at 37 °C for 24 hours prior to analysis.

#### Immunofluorescence microscopy

3T3 or 293ET cells, seeded in 24 well plates on glass coverslips were infected or transfected as described. At the desired time point, cells were washed in PBS and fixed using 3 % paraformaldehyde (PFA, Sigma-Aldrich) in PBS at room temperature for 30 min. After washing in PBS, the cells were quenched with 100 mM NH_4_Cl in PBS and then permeabilised and blocked using 0.1 % Triton with 10 % horse serum (Sigma) or, in the case of experiments including HA staining, 0.1% Saponin with 10 % horse serum in PBS at room temperature for 1 hour. Cells were then incubated with the indicated primary antibodies as detailed in the key resources table. Actin was stained using phalloidin and DNA with stained with 4’6-diamidino-2-phenylindole (DAPI). After several washes in PBS coverslips were then incubated with the associated secondary antibodies. Coverslips were mounted onto slides using Aqua-Poly/Mount (Polysciences, Inc.) and dried overnight at room temperature in the dark.

For transfection experiments, images were captured using a confocal laser scanning microscope (LSM710) (Zeiss GmbH) with a 63x objective. Counting was performed manually, by assessing the actin polymerisation of 100 transfected cells for each cover slip, with 3 coverslips per condition across 2 independent infections. To capture super-resolution images of infected cells, an Olympus CSU-W1 SoRa Spinning Disk was used with 60x lens (Francis Crick Institute). Images were collected with SoRa Disk Changer in Z-stack. Post-image analysis involved running OSR low Macro then deconvolution with SoRa software. All images were processed in Fiji (Image J) and are displayed in colour-blind friendly combinations. To quantify actin foci during in infection, an Axio Imager Upright Microscope (Zeiss) was used. The scorer was blind to which condition was being scored. 3-5 coverslips from 2 independent infections were analysed.

#### Protein sequence analysis

The NCBI Basic Local Alignment Search Tool (BLAST) was used to identify homologues of SteC (UniProt reference D0ZIB5) (https://blast.ncbi.nlm.nih.gov/Blast.cgi). To compare the different SteC protein sequence types among *Salmonella* serovars, 879 complete *Salmonella* genomes were downloaded from Enterobase by searching “Complete Genome” in the “Status” field, which represents the highest assembly quality with circular chromosomes and plasmids (https://enterobase.warwick.ac.uk/, accessed on 2023/06/30). The SISTR1 results from Enterobase were used to identify the subspecies and serovars of the genomes. The *steC* nucleotide sequence from *Salmonella* Typhimurium LT2 (RefSeq: GCF_000006945.2) was used as a reference. A BLAST database was constructed from the *steC* sequence. Each of the 879 *Salmonella* genomes was queried against the *steC* database using BLASTn v2.14.0+ ^29^. The aligned DNA sequences were then extracted and translated into protein sequences using Seqkit v2.4.0 ^30^. The unique SteC protein sequences were summarised and aligned using Clustalo v1.2.4 ^31^. To visualise the SteC types in the *Salmonella* subspecies and serovars, an MStree of the 879 complete *Salmonella* genomes was generated on Enterobase using the rMLST scheme with the MSTree2 algorithm ^32^. The tree was visualised with GrapeTree ^25^. The sequence logo figure for the regions of interest was generated using the ggseqlogo package (https://github.com/omarwagih/ggseqlogo) in R ^33^.

#### Statistical analysis

Statistical significances were calculated using an ordinary one-way analysis of variance (ANOVA) complemented with a *post-hoc* test for multiple comparison’s corrections, as described in figure legends. All analyses were completed on GraphPad Prism (Version 10.1.1).

