## Supplementary figures and tables for "*Salmonella* effector kinase SteC is activated by host-mediated phosphorylation"

This document contains **Figures S1-S5** and **Tables S1-2**.

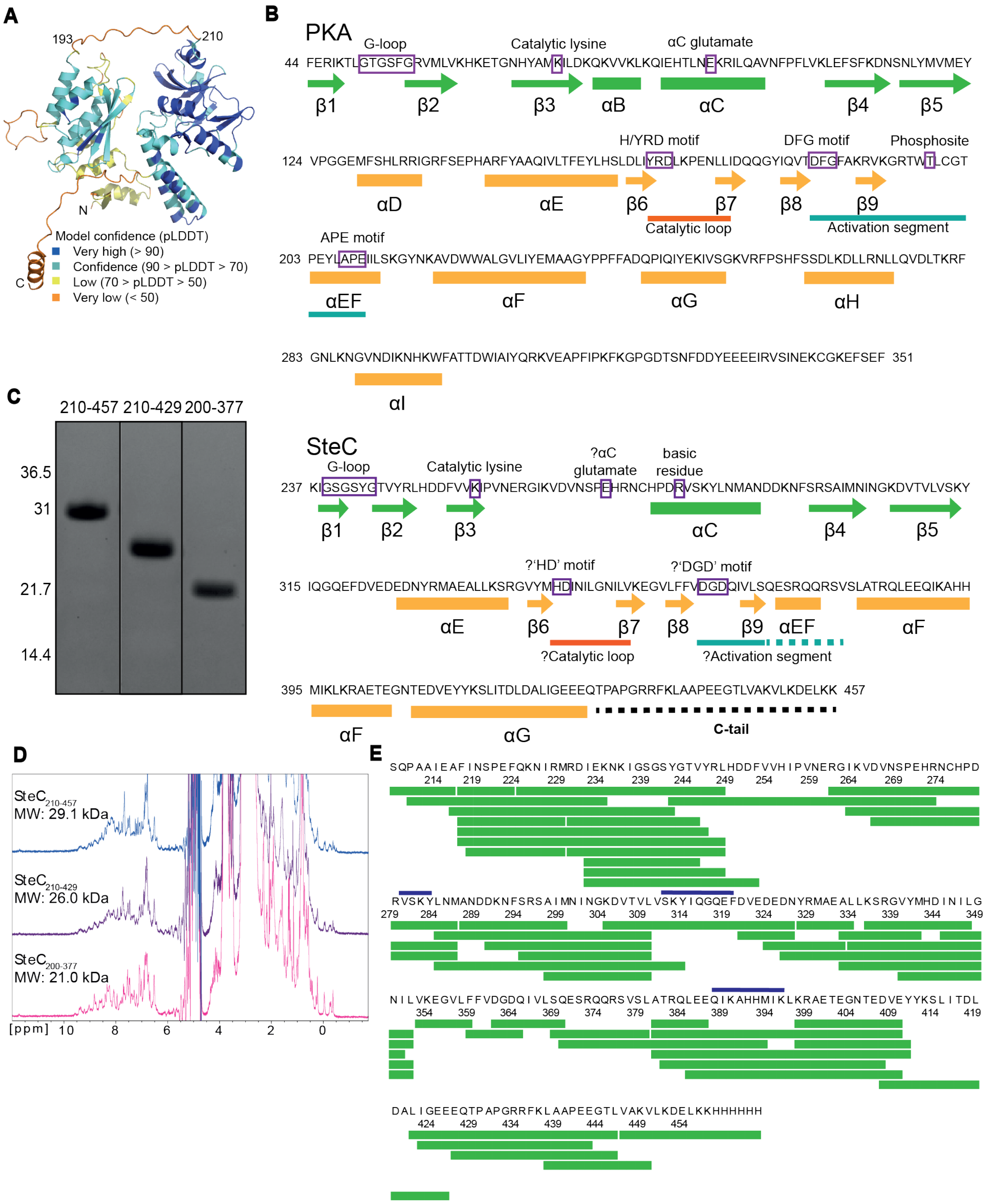


### Figure S1: Predicted and experimentally derived features of the structure of SteC

a) AlphaFold2 prediction of SteC, with N-terminal regulatory domain shown left, kinase domain right, and the C-tail crossing over at the front. Amino acids are coloured with AlphaFold2-generated predicted local distance difference test (pLDDT) score as per key.

b) Top: PKA key sequence motifs (purple) are highlighted alongside secondary structure elements: α-helices and β-sheets comprising the N lobe (green) and C lobe (orange); the catalytic loop, activation segment and activation loop are indicated. Adapted from Sheetz and Lemmon (2022)[26]. Bottom: SteC with secondary structure elements as per AF prediction in N lobe (green) and C lobe (orange); hypothetical key motifs, and possible catalytic loop, activation segment, activation loop and C-tail are indicated in purple, with dotted lines referring to non-canonical features.

c) SDS-PAGE analysis of purified SteC encompassing amino acids 210-457, 210-429 and 200-377.

d) 1D $^{1}$H-NMR spectra of SteC K256H mutants of 210-457, 210-429 and 200-377 expressed in *E. coli* collected at 25 °C on a Bruker AVANCE spectrometer operating at 800 MHz.

e) Peptide coverage of Hydrogen deuterium exchange experiments of SteC 210-457 K256H. Total 53 peptides, 100% coverage, 3.80 redundancy. Regions significantly protected in the presence of the FMNL1 peptide are highlighted in blue above. Regions of SteC demonstrating protection in the presence of the peptide (at least 0.45 difference in all peptides covering this region) are highlighted on the AF predicted structure in blue in **Figure 1c**.

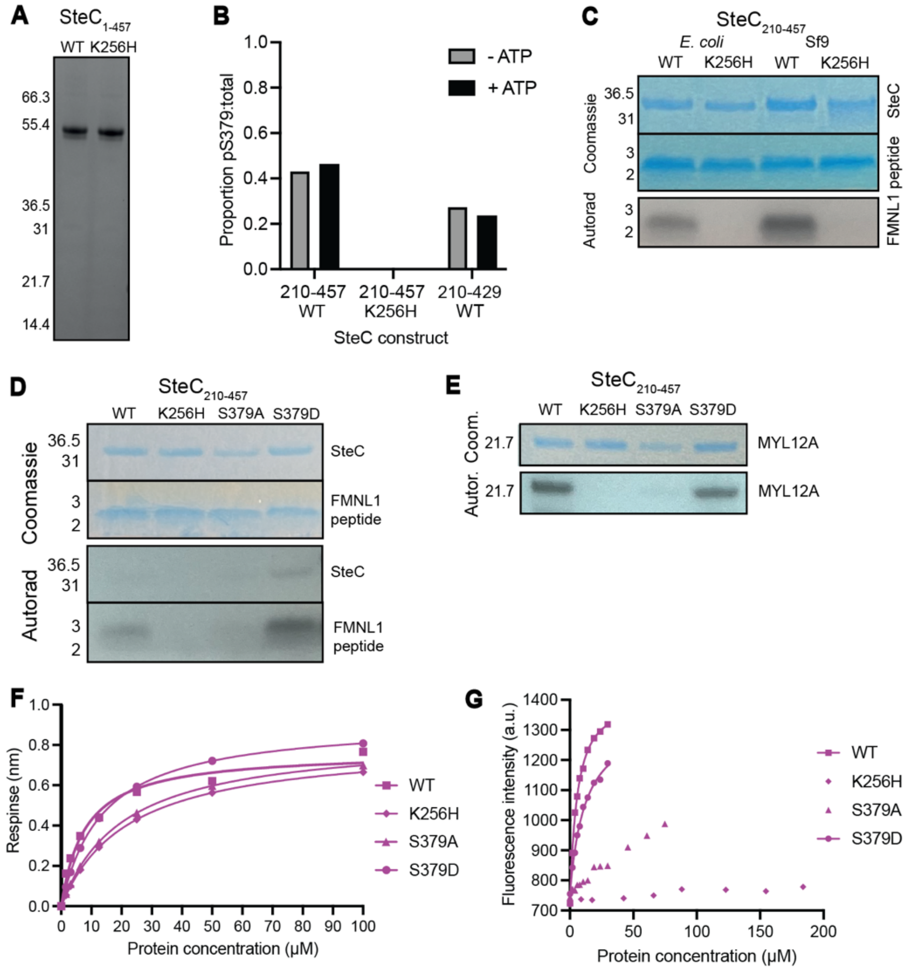


**Figure S2: Kinase activity and substrate binding of SteC**

a) Coomassie stained SDS-PAGE analysis SteC$_{1-457}$ WT and K256H expressed in Sf9 cells.

b) *E. coli* expressed SteC, either 210-457 or 210-429, was incubated with or without ATP and analysed by mass spectrometry for the phosphorylation of the S379 containing peptide. Experiment performed once.

c) Radioactive kinase assay comparing insect cell and *E. coli* purified SteC against FMNL1 peptide. Quantification of three repeats is shown in **Figure 2c**, with pairwise statistical comparisons as above.

d) Radioactive kinase assay of SteC$_{210-457}$ WT and mutants expressed in *E. coli* at 5 µM with 100 µM FMNL1 peptide. The assay is representative of 3 repeats.

e) Radioactive kinase assay of 100 nM SteC$_{210-457}$ WT and mutants expressed in *E. coli* with 5 µM MYL12A. The assay is representative of 3 repeats.

f) Analysis of the interaction of $SteC_{210\text{-}429}$ WT and mutants with an FMNL1 peptide by Biolayer Interferometry. The biotinylated peptide was loaded onto Streptavidin sensors. Fitted curves are shown as purple lines.

g) Fluorescence titrations of 500 nM mant-AMPPNP with $SteC_{210\text{-}429}$ WT and mutants expressed in *E. coli*, as in **Figure 2g**. Higher concentrations of $SteC_{K256H}$ and $SteC_{S379A}$ were used. Fitted curves are shown as purple lines.

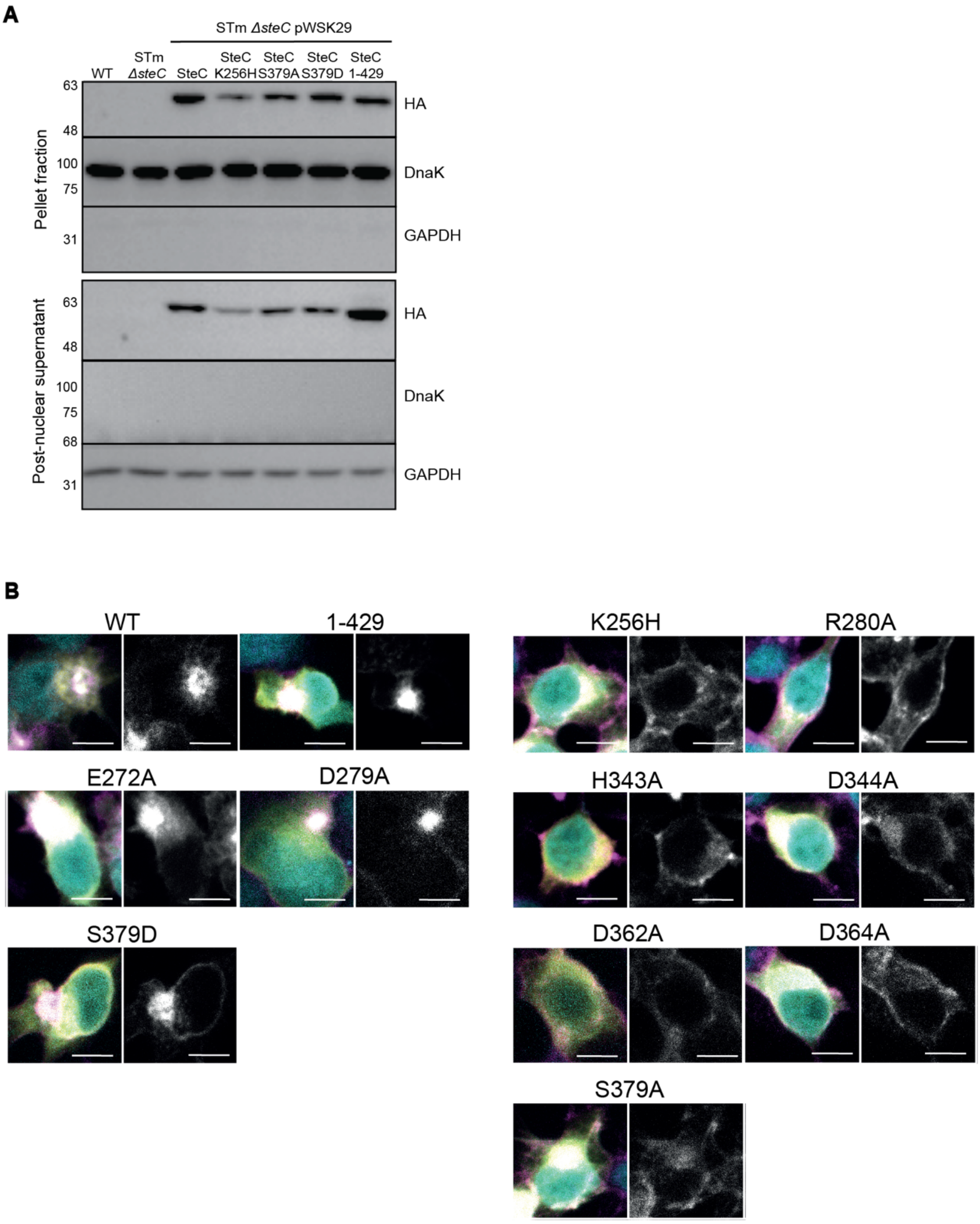


**Figure S3: SteC in infected and transfected mammalian cells**

a) 3T3 cells were infected with *Salmonella* WT, *ΔsteC* and *ΔsteC psteC* strains. Eight hours post invasion, cells were lysed and pellet and post-nuclear fractions analysed by immunoblotting with antibodies against HA (SteC), DnaK and GAPDH. Data representative of 2 independent biological repeats.

b) HEK 293ETs were transfected with the indicated GFP-SteC variants and following fixation, were stained with 4’6-diamidino-2-phenylindole (DAPI) and phalloidin AF647 prior to imaging. Representative merged images with DAPI (cyan), phalloidin (magenta) and GFP (yellow) and a grayscale for the phalloidin channel (right) are shown for each condition. Representative cells were chosen randomly. Scale bar represents 5 μm.

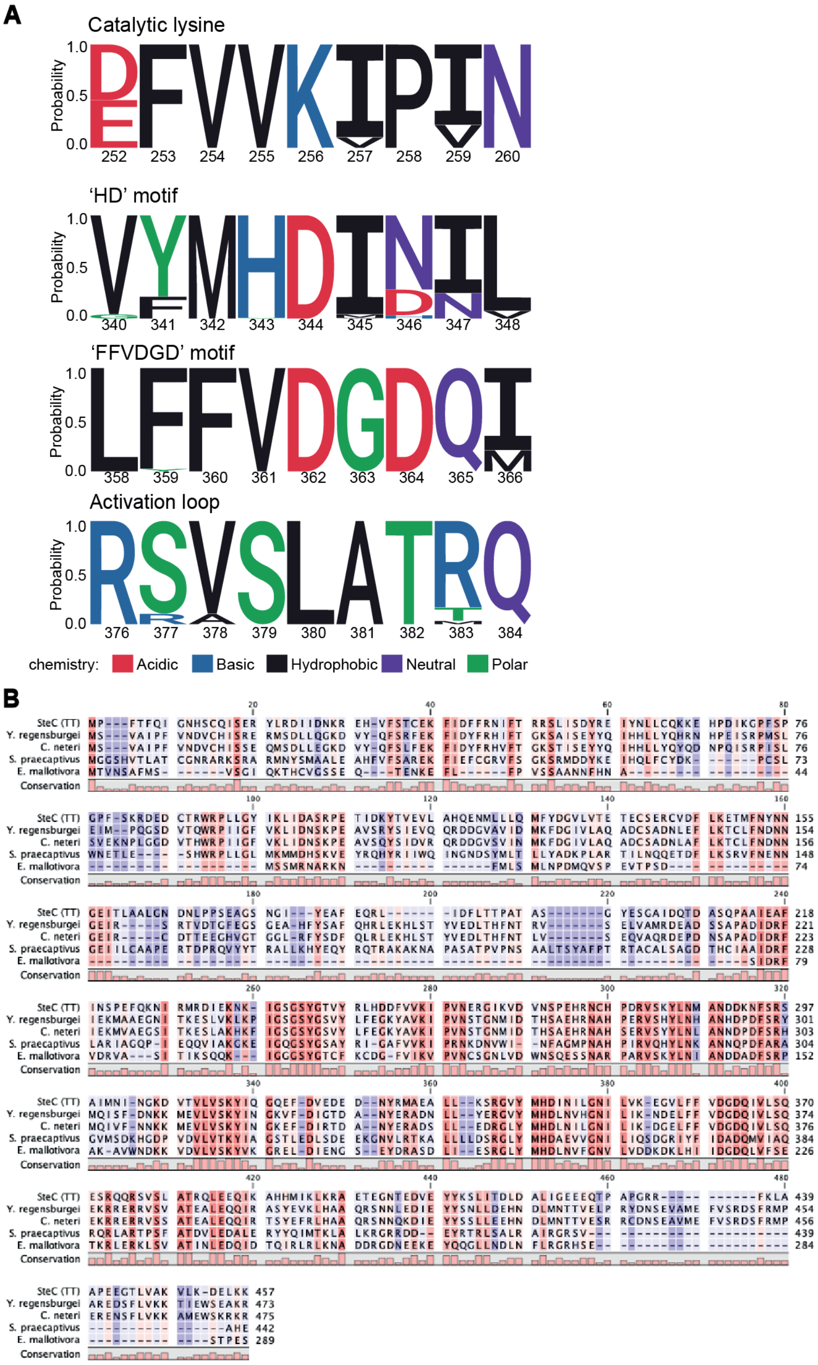


**Figure S4: Sequence conservation across putative homologs of SteC**

a) Regions of interest were compared in the same 879 genomes from *Salmonella* clinical isolates analysed in **Figure 4a**. Sequence logos for these motifs were generated using the ggseqlogo package in R[35].

b) A multiple sequence alignment was performed of the amino acid sequences of SteC and its top four bacterial homologs identified by BLASTp search: *Yokenella* *regensburgei* (98% coverage, 41% amino acid identity), *Cedecea neteri* (92%, 42%), *Sodalis praecaptivus* (86%, 34%) and *Erwinia mallotivora* (42%, 49%) using T-Coffee in CLC Sequence Viewer 7. The background colour scale represents a spectrum from full homology (red) to complete lack of homology (light blue). The conservation graph below demonstrates the number of amino acid sequences sharing any given residue.

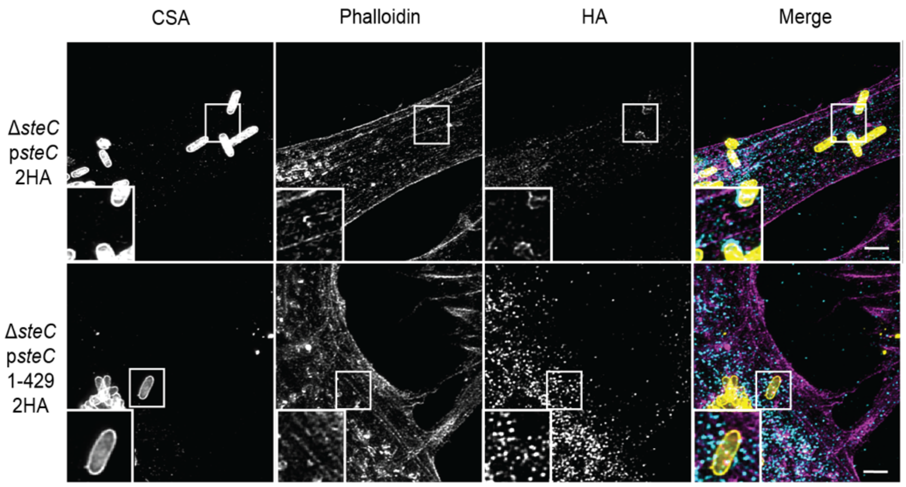


**Figure S5: SteC associated with *Salmonella* in infected cells**

Second set of representative images from super resolution microscopy analysing subcellular localisation of HA-tagged SteC variants following the infection of 3T3 cells, as shown in **Figure 4b**. CSA labels *Salmonella* and Phalloidin stains actin. In the merge, CSA is shown in yellow, phalloidin in magenta and HA in cyan. Scale bars represent 5 μm.

### Supplementary Tables

#### Table S1: Phosphorylated peptides of FMNL1 after incubation with SteC and ATP

$FMNL1_{1\text{-}458}$ and $SteC_{1\text{-}457}$ were expressed in Sf9 cells. FMNL1 was analysed by phospho-MS either alone or after incubation with SteC and ATP. No FMNL1 phosphosites were detected in the control condition. Phosphorylated FMNL1 peptides after incubation with SteC and ATP are reported here. Data were analysed with MaxQuant[28].

| Starting amino acid | Peptide sequence | Mass (Da) | Phosphorylation site |
|---|---|---|---|
| 178 | NKPLEQ**pS**VEDLSK | 1565.73 | S185 |
| 178 | NKPLEQ**pS**VEDLSKGPPSSVPK | 2315.14 | S185 |
| 191 | GPPSSVPK**pS**R | 1090.56 | S199 |
| 199 | SRHL**pT**IK | 933.48 | T203 |
| 199 | SRHL**pT**IKLTPAHSR | 1695.89 | T203 |
| 201 | HL**pT**IKLTPAHSRK | 1580.86 | T203 |
| 201 | HL**pT**IKL**pT**PAKSR | 1532.73 | T203 & T207 |

**Table S2: Phosphorylated and unphosphorylated peptides of recombinant SteC with and without ATP**

$SteC_{1-457}$ was expressed in Sf9 cells and was analysed either alone or after incubation with ATP. S75 was barely detected prior to the addition of ATP and is low-medium phosphorylated on the addition of ATP. S379 was highly phosphorylated in both conditions. Peptide intensity is reported in a logarithmic scale. Data were analysed with MaxQuant.

| **Starting amino acid** | **Peptide sequence** | **Phosphorylation site** | **Intensity (-ATP)** | **Intensity (+ATP)** |
|---|---|---|---|---|
| 66 | EHPDIKGPFSPGPFSK | | 6.52263 | 7.67358 |
| 66 | EHPDIKGPF**pS**PGPFSK | S75 | NaN | 7.25220 |
| 72 | GPFSPGPFSK | | 8.92893 | 8.82299 |
| 72 | GPF**pS**PGPFSK | S75 | 7.20550 | 8.09496 |
| 377 | SVSLATR | | 7.60412 | 7.28526 |
| 377 | SV**pS**LATR | S379 | 9.35727 | 9.25940 |
